# SLD5/GINS4 controls dynein-dependent centrosome maturation and exposes a candidate mitotic vulnerability in cancer

**DOI:** 10.64898/2026.05.07.723511

**Authors:** Vipin Kumar, Vivek Singh, Raksha Singh, Praveen Kumar, Tanushree Ghosh

## Abstract

Faithful proliferation requires coordinated DNA replication with centrosome maturation and spindle-pole integrity. SLD5, encoded by GINS4, is a core component of the GINS replication complex and is frequently elevated in tumors, but whether it links replication-associated cancer states to centrosome control has remained unclear. Here, we show that GINS4/SLD5 is recurrently upregulated across human cancers at transcript and protein levels and marks tumor programs enriched for DNA replication, chromosome segregation, and mitotic control. In cancer cells, Sld5 depletion dispersed PCM1, AZI1, and CEP290-positive centriolar satellites without eliminating these satellite proteins, reduced dynein heavy chain expression, and destabilized dynein-dynactin localization at spindle poles. Direct depletion of dynein heavy chain, co-depletion analyses, and pharmacological inhibition of dynein motor activity with ciliobrevin D phenocopied Sld5 loss, causing satellite dispersion, defective recruitment of PLK1, Aurora A, CEP192, and CEP215 to centrosomes, and multipolar spindle formation. These defects occurred without detectable DNA damage or checkpoint activation, indicating a non-canonical Sld5 function beyond its role in the replisome. Cancer dependency and kinase network analyses further nominate SLD5-associated mitotic and checkpoint pathways as therapeutic targets. Our findings identify SLD5/GINS4 as a regulator of dynein-dependent centrosome maturation and a candidate vulnerability in replication-driven cancers, with potential value for biomarker-guided therapeutic stratification.

**Graphical abstract:** 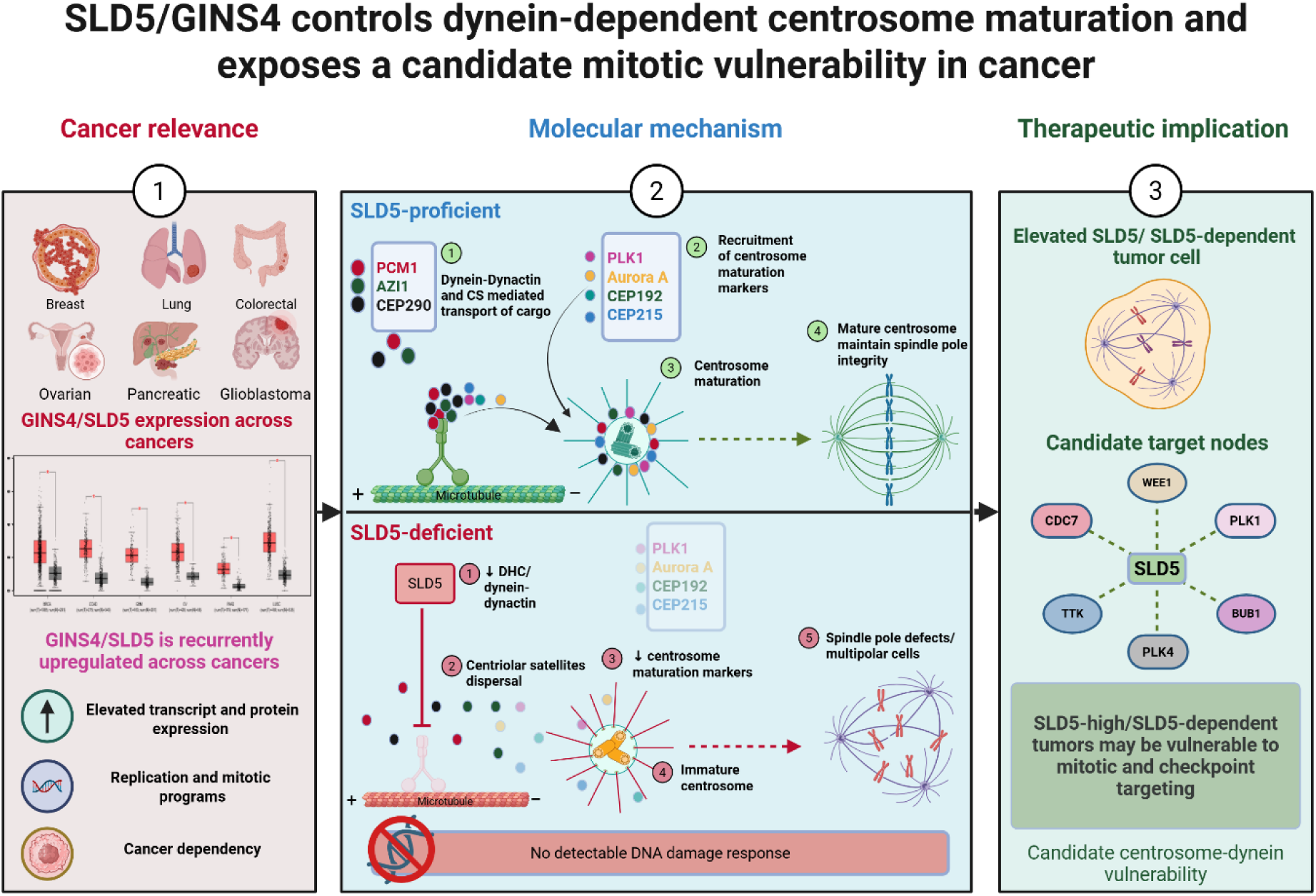

## Introduction

Faithful cell division requires coordination between DNA replication, centrosome maturation, and mitotic spindle assembly. The centrosome is the principal microtubule-organizing center of most animal cells and is built from a pair of centrioles embedded in the pericentriolar material (PCM), a dynamic protein matrix that nucleates and anchors microtubules. During cell-cycle progression, PCM content is relatively limited after mitosis but expands sharply during late G2 and early mitosis, increasing the ability of centrosomes to recruit γ-tubulin ring complexes and organize robust bipolar spindle poles. This process, termed centrosome maturation, is therefore essential for chromosome segregation, mitotic fidelity, and genome stability [1–3]. Centrosome maturation is driven by a coordinated kinase-scaffold network. CEP192 acts as an organizing platform that recruits and activates Aurora A and polo-like kinase 1 (PLK1) at centrosomes, initiating a phosphorylation cascade that promotes PCM expansion and bipolar spindle assembly [4]. PLK1- and Aurora A-dependent phosphorylation of centrosomal substrates, including pericentrin, CEP215/CDK5RAP2, and TACC-family proteins, supports PCM scaffold assembly and enhances microtubule-nucleating activity [5, 6]. Perturbation of this program weakens spindle-pole architecture and can promote chromosome missegregation, chromosomal instability, and centrosome-associated mitotic vulnerability, features that are particularly relevant in cancer [7–9]. Centriolar satellites are important but incompletely understood regulators of centrosome maintenance. These non-membranous, PCM1-positive granules cluster around centrosomes and move dynamically along microtubules towards the pericentrosomal region. PCM1 was originally characterized as a cell-cycle-regulated centrosome-associated protein and was subsequently established as a core scaffold required for centriolar satellite assembly and organization [10–13]. Beyond PCM1, centriolar satellites contain numerous proteins that also localize to centrosomes or cilia and contribute to centrosome organization, ciliogenesis, protein trafficking, and centrosome proteostasis [14–16]. The pericentrosomal positioning of centriolar satellites depends on microtubules and minus-end-directed transport by cytoplasmic dynein. Dynein is a multi-subunit motor complex in which the dynein heavy chain contains the AAA+ ATPase motor domain that powers movement along microtubules, whereas intermediate, light-intermediate, and light chains contribute to complex stability, regulation, and cargo linkage [17, 18]. Dynein acts together with dynactin, a cargo-adaptor and processivity-enhancing complex that is required for efficient dynein-mediated transport[19–22]. Consistent with this model, perturbation of microtubules or of the dynein-dynactin complex disperses PCM1-positive satellites and reduces centrosomal recruitment of proteins such as centrin, ninein, and pericentrin [12]. Satellite-associated cargoes, including CEP290, CEP72, CEP131/AZI1, and related centrosomal modules, further support the idea that centriolar satellites function as transport and storage hubs for centrosome-regulatory proteins [23–26]. However, how dynein-dependent satellite transport is coupled to centrosome maturation in proliferating cancer cells remains insufficiently defined. SLD5, encoded by GINS4, is classically recognized as a component of the GINS complex, which associates with CDC45 and the MCM2-7 helicase to form the active CMG helicase required for DNA replication initiation and fork progression [27–30]. Because malignant cells frequently experience high proliferative demand and replication stress, replication-associated genes are often activated in tumors. Consistent with this, GINS4 has been reported to be upregulated in multiple cancers and has been linked to tumor growth, poor prognosis, and immune-related tumor states in both tumor-specific and pan-cancer analyses [31–35]. These observations have largely framed GINS4 as a replication-associated oncogenic factor, but they do not fully explain how SLD5 supports cancer-cell fitness during mitotic progression. A non-canonical centrosomal function of SLD5 was suggested by previous work showing that SLD5 localizes to centrosomes and preserves centriolar satellites during chromosome congression, thereby protecting centrosomes from mitotic forces[36]. This observation raises the possibility that SLD5 connects replication-associated proliferative states to the dynein-dependent trafficking machinery required for centrosome maturation. In this study, we integrate pan-cancer profiling with mechanistic cell-biology analyses to define how SLD5 regulates centriolar satellite organization, dynein-dynactin function, centrosome maturation, and spindle-pole integrity. We further evaluate whether the SLD5-dynein-centrosome axis represents a therapeutically exploitable vulnerability in GINS4-high and SLD5-dependent cancers.

## Methods

### Pan-cancer GINS4/SLD5 expression analysis

Pan-cancer GINS4/SLD5 mRNA expression was evaluated using publicly available tumor transcriptomic datasets from TCGA and corresponding normal tissue datasets from GTEx, accessed via cancer genomics portals such as GEPIA and UCSC Xena. Expression values were analyzed as log-transformed TPM-normalized data where applicable, and tumor versus normal comparisons were performed across cancer types using platform-reported statistical outputs or standard group-wise comparisons. Cancer-type abbreviations followed the TCGA nomenclature [37–40].

### Pan-cancer protein expression and immunohistochemistry analysis

GINS4 protein abundance across tumor groups was analyzed using public pan-cancer proteomic resources, including TCPA/TPCPA-derived tumor proteomic datasets. Protein-level differences among tumor lineages were assessed using group-wise comparisons and one-way ANOVA, as appropriate. Representative immunohistochemistry images and staining annotations were obtained from the Human Protein Atlas, and semi-quantitative H-score comparisons were used to estimate relative GINS4 protein expression between normal and tumor tissues [41–45].

### Integrated Stouffer’s Z-score analysis

To generate an integrated pan-cancer ranking, cancer-type-level statistics were converted into signed Z-scores and combined using Stouffer’s method. Genes were ranked according to their combined Stouffer’s Z-score, and GINS4/SLD5 was highlighted within the ranked distribution. Cancer-specific Z-scores were visualized for tumor types showing positive GINS4/SLD5 enrichment [46, 47].

### Immune infiltration, copy-number, and survival analyses

Associations between GINS4/SLD5 expression and tumor immune infiltration were assessed using TIMER/TIMER2-based immune deconvolution outputs. Immune-cell compartments included B cells, CD8⁺ T cells, CD4⁺ T cells, macrophages, neutrophils, and dendritic cells. Copy-number-associated immune infiltration patterns were analyzed using TIMER copy-number categories, including deep deletion, arm-level deletion, diploid/normal, arm-level gain, and high amplification. Survival associations were evaluated using Kaplan–Meier and Cox proportional hazards analyses from public cancer survival platforms, and log-rank P values or hazard ratios were used to summarize outcome associations [48–50].

### GINS4/SLD5-associated gene correlation and pathway enrichment

Genes associated with GINS4/SLD5 expression were identified using Pearson or Spearman correlation analysis across tumor datasets. Ranked gene lists were subjected to functional enrichment analysis using Gene Ontology, MSigDB, KEGG, and Reactome pathway collections. Enrichment results were summarized using normalized enrichment scores and FDR-adjusted P values. Enrichment-map and UpSet-style plots were used to visualize pathway clustering and overlap among enriched cellular-component annotations [51–57].

### Cancer dependency and SLD5–DYNC1H1 association analysis

Cancer cell line dependency data were obtained from DepMap/CCLE resources. SLD5/GINS4 loss-of-function effects were evaluated using CRISPR-Cas9 dependency or log fold-change values, and cancer-lineage-specific distributions were generated to identify tumor contexts with increased sensitivity to SLD5 perturbation. Associations between SLD5/GINS4 and DYNC1H1 were assessed using matched expression and dependency datasets, with Pearson and Spearman correlations calculated within each cancer lineage [58–60].

### GINS4/SLD5–POLR2A correlation and structural prediction

The relationship between GINS4/SLD5 and POLR2A expressions was examined across pan-cancer datasets using correlation analysis. Cancer-type-specific Pearson and Spearman coefficients were calculated. A predicted GINS4–POLR2A protein-complex model was generated using AlphaFold/ColabFold-based structure prediction, and confidence was assessed using predicted TM-score, interface predicted TM-score, and predicted aligned error outputs [61, 62].

### Kinase enrichment, therapeutic network, and co-occurrence analysis

To identify candidate therapeutic nodes associated with the SLD5/GINS4 program, GINS4-associated genes were analyzed using kinase enrichment approaches. Candidate kinases were ranked by enrichment score or Z-score, and selected mitotic, replication, and checkpoint-associated kinases. Survival relevance of candidate kinases was assessed using hazard ratios from public cancer survival datasets. Genetic co-occurrence among GINS4/SLD5 and candidate kinase genes was evaluated using cBioPortal, and significant co-occurrence relationships were summarized [63–65].

### Immunofluorescence

For indirect immunofluorescence to visualize α-tubulin, γ-tubulin, Plk1, and Aurora-A, HeLa cells were grown on coverslips and fixed with 4% formaldehyde for 10 min, then with ice-cold methanol for 5 min. For visualization of Sld5, the cells were pre-permeabilized in the extraction buffer (80 mM PIPES, 1 mM MgCl2, 1 mM EGTA, and 0.5% Triton X-100) for 2 min, followed by fixation with ice-cold methanol for 2 min. To visualize Cep215, cells were fixed with ice-cold methanol for 10 min. Samples were blocked with 3% BSA or 10% fetal bovine serum (FBS) for 30 min, then stained with the specific primary antibody (supplementary table S1) for 1 h, followed by incubation with the secondary antibody for 1 h. Finally, the cells were visualized under a microscope after mounting with Vectashield containing DAPI, which stains the nucleus. The secondary antibodies used were Alexa Fluor 488- or Alexa Fluor 555-conjugated and purchased from Invitrogen. We assayed the intensity of the Alexa Fluor 488 or Alexa Fluor 555 signal using the ImageJ software (NIH) to quantify the levels of Cep215, Plk1, Aurora-A, and γ-tubulin signals at the spindle poles. The mean intensity of Alexa Fluor signal for spindle poles was marked as low intensity if the average mean intensity was less than 50% of the average mean intensity of the signal observed in control transfected cells.

### Immunoprecipitation

Experimental cells were harvested, washed with PBS, and lysed with TNN buffer (50 mM Tris, pH 8.0, 120 mM NaCl, 1% NP40 & protease inhibitor) for 30 min on ice. The lysate was centrifuged at 20,000 rcf for 15 min at 4 °C. The supernatant was collected, and protein was quantified using a BCA assay kit (BioRad) on a SYNERGY microplate reader. An equal amount of protein was used for the immunoprecipitation assay. To remove nonspecific binding, the supernatant was pretreated with protein A/G agarose (Abmart, Cat. # A10001M) and IgG (Santa Cruz #sc-2025) for 1 h at 4 °C. After incubation and centrifugation, the supernatant was collected into a new tube, and the supernatant was incubated with the indicated antibodies (supplementary table S2) overnight. Further, samples were incubated with 30 μL of protein A/G agarose for 2 h at 4 °C. Afterward, all samples were washed 5 times with TNN buffer, eluted with 2x sample buffer, and boiled for 10 min at 95 °C. Interactions were analyzed by western blotting with specific antibodies, and whole-cell lysate was used as a control (input).

### Western blotting

For Western blotting, whole-cell lysates from cells at almost equal confluence were prepared in a 1:1 volume ratio with Laemmli buffer and denatured at 95°C, followed by sodium dodecyl sulfate-polyacrylamide gel electrophoresis (SDS-PAGE). The gel was transferred to a nitrocellulose membrane and blocked with 3% bovine serum albumin (BSA) in 1× Tris-buffered saline with Tween 20 (TBST). The membrane was then incubated with the appropriate antibodies (supplementary table S3), washed, and probed with horseradish peroxidase (HRP)-conjugated secondary antibody. Enhanced chemiluminescence was used to visualize the protein bands.

### Antibodies

Sld5 antibody, raised against amino acids 1-198 of human Sld5, used for Western blotting and immunofluorescence assays, was procured from Abcam (Ab101346). For Western blotting, the following antibodies were used: Plk1, AZI1, Cep290, DYNC1I1, DYNC1I2, DYNC1LI1, DYNC1LI2, DYNLT1, DYNLT3, DYNLL1, MCM5, RNA polymerase II, and BubR1, all purchased from Abcam. Anti-DYNC1H1 antibody was obtained from Millipore. Anti-β-actin antibody was obtained from Santa Cruz Biotechnology. Anti-Aurora-A antibody was procured from Cell Signaling Technology. For indirect immunofluorescence assays, the following antibodies were used: Antibodies against-α-tubulin, γ-tubulin, Plk1, Cep215, DYNLT1, and DCTN2 (p50) were obtained from Abcam. Anti-Aurora-A and anti-PCM1 antibodies were procured from Cell Signaling Technology. An anti-Cep192 antibody was procured from Novus. Anti-DYNC1H1 mouse antibody was procured from Sigma. Secondary antibodies used were Alexa Fluor 488- and Alexa Fluor 555-conjugated and were purchased from Invitrogen. For immunoprecipitation, the following antibodies were used: an Sld5 antibody from Abcam and a γ-tubulin antibody from Cell Signaling Technology.

### Cell culture and drug treatment

The cell line was maintained at 37°C in Dulbecco’s modified Eagle’s medium (DMEM) supplemented with 10% FBS and 1% antibiotic and anti-mycotic solution. HeLa cells were treated with 10 µM MG132 for 90 min, and subsequently, 50 µM ciliobrevin D (Sigma) was dissolved in dimethyl sulfoxide (DMSO) (Sigma).

### RNAi silencing, and reverse-transcriptase PCR

Transfection of siRNA targeting the endogenous genes was performed using Lipofectamine 2000 (Invitrogen). Cells were transfected with specific siRNA duplexes (40–80 nM) on three consecutive days and harvested 24 h after the last transfection for immunoblotting, immunofluorescence, or reverse-transcriptase PCR. Microtubule nucleation assay, HeLa cells were transfected with three consecutive days then treated with 100 nM nocodazole to destabilization of MT and then released for 5 h to make new MT and CS. For reverse transcriptase PCR, RNA was isolated using the standard TRIzol method, and 1 µg of RNA was used for cDNA synthesis. Details of the protocols and DNA primers (supplementary table S4) used are available on request. The siRNA sequences used for targeting the human genes were as follows: GL2, CGUACGCGGAAUACUUCGA; SLD5 (1), AAAUGGAGAUGGAGAGGAU; SLD5 (2), GCUGGAGAGCAAGCCUGAGAUUGUA; KID, AAGAUUGGAGCUACUCGUCGU; BUBR1, AACGGGCAUUUGAAUAUGAAA; MAD2, GCGUGGCAUAUAUCCAUCU; DHC, GAGAGGAGGUUAUGUUUAAUU.

### Statistical analysis and visualization

Unless otherwise stated, analyses were performed using standard statistical approaches in R, Python, or web-based bioinformatics platforms. Continuous variables were compared using parametric or nonparametric tests, depending on the data distribution. Correlations were assessed using Pearson or Spearman coefficients. Survival outcomes were analyzed using Kaplan–Meier curves, log-rank tests, and Cox proportional hazards models. Multiple testing was controlled using FDR adjustment, where applicable, and P < 0.05 or FDR < 0.05 was considered statistically significant.

## Results

### GINS4 is recurrently upregulated across human cancers at the transcript and protein levels

To define the cancer-wide expression landscape of GINS4, we first analyzed its mRNA abundance across tumor and matched or corresponding normal tissues. GINS4 was significantly elevated in multiple tumor types, indicating that its transcriptional activation is a recurrent feature of human malignancy rather than a cancer-type-restricted event (Fig. 1A and Supplementary Table 5). Consistent with the transcriptomic data, tumor proteomic profiling showed marked variation in GINS4 protein abundance across cancer groups, with a highly significant difference among tumor lineages by one-way ANOVA (P = 9.22 × 10⁻¹⁸; Fig. 1B). These data support a broad pan-cancer increase in GINS4 at both RNA and protein levels. We next evaluated GINS4 protein expression in representative tissue sections using immunohistochemistry. Compared with normal liver/hepatocytes, liver cholangiocarcinoma showed stronger GINS4 staining, with an approximate H-score increase from 75 to 200, corresponding to a 2.67-fold increase. Similarly, prostate adenocarcinoma showed stronger GINS4 staining relative to the normal comparator tissue, with an approximate H-score increase from 130 to 255, corresponding to a 1.96-fold increase (Fig. 1C). Thus, independent protein-level evidence confirms enhanced GINS4 expression in tumor tissue. To integrate the pan-cancer signal at the gene-ranking level, we calculated genome-wide Stouffer’s Z-scores. GINS4, also referred to as SLD5, was positioned within the positive tail of the ranked distribution, with a Stouffer’s Z-score of 9.14 (Fig. 1D and Supplementary Table 5). Cancer-specific Z-score analysis further confirmed positive GINS4 enrichment across multiple malignancies, including ACC, KICH, KIRC, KIRP, LGG, LIHC, LUAD, MESO, PAAD, PCPG, PRAD, SARC, THCA, and UCEC, with the strongest signals observed in LGG, MESO, and KIRP (Fig. 1E). Collectively, these findings identify GINS4 as a recurrently upregulated pan-cancer gene with consistent transcriptomic, proteomic, and histological evidence of tumor-associated activation.

**Fig. 1.**
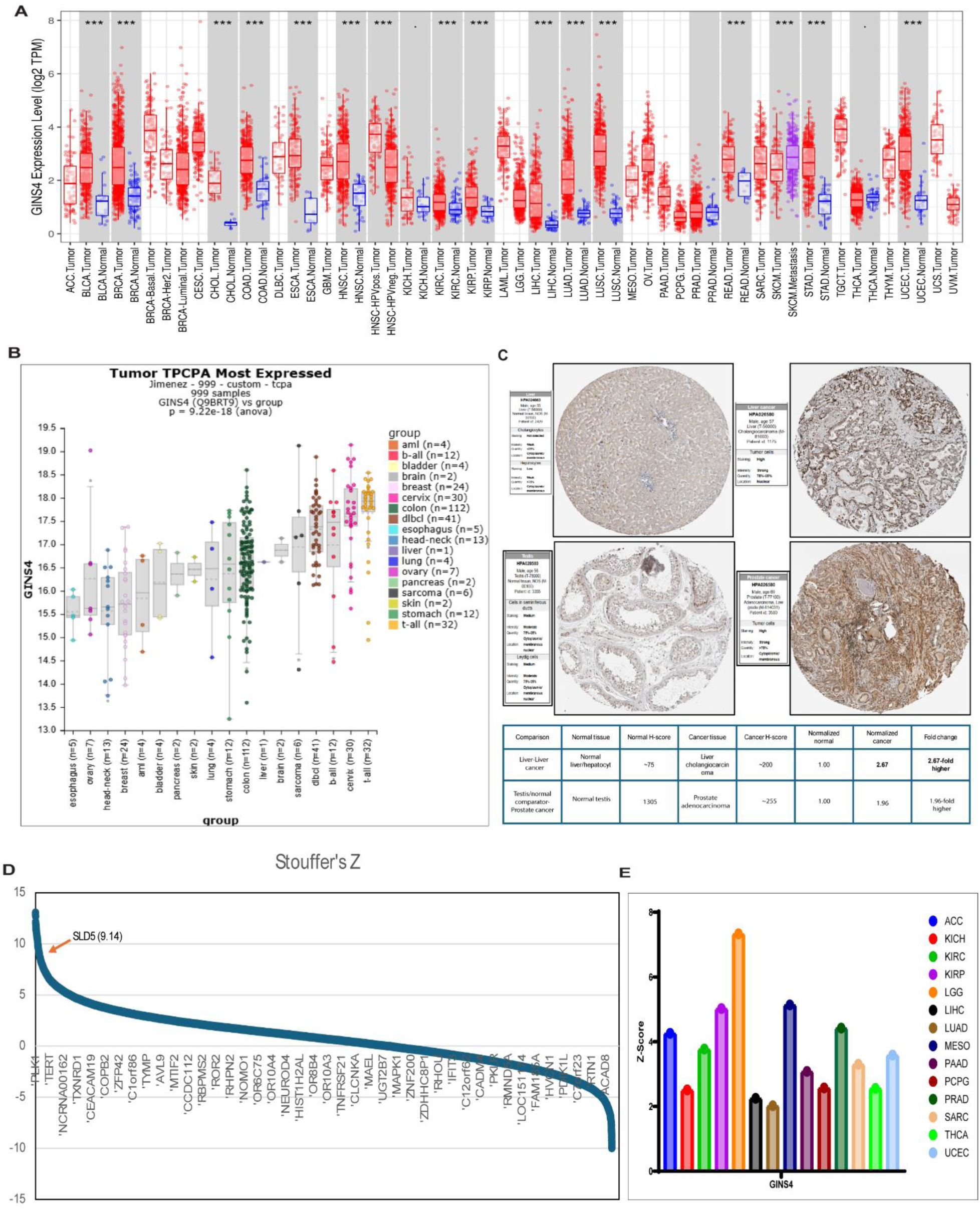
Pan-cancer expression landscape and integrated ranking of GINS4. **A.** Pan-cancer analysis of GINS4 mRNA expression across tumor and normal tissues. Expression is shown as log₂ TPM. Red points denote tumor samples and blue points denote normal samples; additional sample classes are shown where indicated in the plot. Box plots show the median and interquartile range, with whiskers indicating the distribution range. Statistical significance for tumor vs. normal comparisons is indicated above each type of cancer. **B.** Tumor proteomic analysis of GINS4 across cancer groups from the TPCPA/TCPA proteomic dataset. Each point represents an individual tumor sample, and box plots summarize GINS4 protein abundance within each tumor group. Group-level differences were assessed by one-way ANOVA; P = 9.22 × 10⁻¹⁸. **C.** Representative immunohistochemistry images showing GINS4 protein staining in normal and tumor tissues. GINS4 staining increased in liver cholangiocarcinoma compared with normal liver/hepatocytes, with an approximate H-score increase from 75 to 200, corresponding to a 2.67-fold increase. GINS4 staining was also increased in prostate adenocarcinoma compared with the indicated normal comparator tissue, with an approximate H-score increase from 130 to 255, corresponding to a 1.96-fold increase. **D.** Genome-wide Stouffer’s Z-score distribution from the integrated pan-cancer analysis. Genes were ranked from the highest to the lowest Stouffer’s Z-score. GINS4/SLD5 is highlighted in the positive tail of the distribution with a Stouffer’s Z-score of 9.14. **E.** Cancer-specific Stouffer’s Z-scores for GINS4 across selected tumor types. Positive Z-scores indicate recurrent enrichment of GINS4-associated signal in the indicated cancer type. ACC, adrenocortical carcinoma; KICH, kidney chromophobe; KIRC, kidney renal clear cell carcinoma; KIRP, kidney renal papillary cell carcinoma; LGG, lower-grade glioma; LIHC, liver hepatocellular carcinoma; LUAD, lung adenocarcinoma; MESO, mesothelioma; PAAD, pancreatic adenocarcinoma; PCPG, pheochromocytoma and paraganglioma; PRAD, prostate adenocarcinoma; SARC, sarcoma; THCA, thyroid carcinoma; UCEC, uterine corpus endometrial carcinoma.

### GINS4-associated tumor states are linked to immune context and replication cell-cycle programs

To determine whether GINS4 expression is associated with the tumor microenvironment, we assessed immune infiltration patterns across selected cancer types. GINS4 expression showed cancer-specific relationships with tumor purity and immune-cell infiltration, including B cells, CD8⁺ T cells, CD4⁺ T cells, macrophages, neutrophils, and dendritic cells (Supplementary Fig. S1A). Kaplan–Meier analyses further indicated that GINS4 expression and immune-infiltration strata were associated with survival differences in selected tumor contexts, suggesting that the biological impact of GINS4 may depend on lineage-specific tumor-immune states (Supplementary Fig. S1B). Copy-number-based analyses also revealed that GINS4 genomic alterations were associated with variation in immune infiltration levels across tumor types (Supplementary Fig. S1C). Because GINS4 is a component of DNA replication machinery, we next examined the functional programs associated with GINS4-linked transcriptional states. Gene set enrichment analysis revealed strong positive enrichment for DNA replication, chromosome segregation, DNA recombination, double-strand break repair, spindle organization, regulation of the mitotic cell cycle, cytokinesis, sister chromatid cohesion, and regulation of DNA repair (Supplementary Fig. S1D-F). In contrast, negatively enriched or lower-scoring programs included extracellular structure organization, cell–cell adhesion, regulation of locomotion, and tissue/vascular-associated processes. Cellular-component overlap analysis further showed enrichment of nuclear, nucleoplasmic, chromatin, chromosome, MCM complex, DNA replication factor A complex, GINS complex, and spindle-pole annotations (Supplementary Fig. S1G). These analyses indicate that GINS4-high tumor states are characterized by replication, DNA repair, and mitotic cell-cycle programs. After this, we explored more cellular functions of the SLD5 in cancer cell lines.

### Sld5 maintains centriolar satellite organization and spindle-pole integrity

Centriolar satellites are pericentrosomal protein assemblies that support centrosome organization by acting as reservoirs for centrosomal factors. We previously reported that loss of Sld5 leads to the dissipation of PCM1, a core scaffolding component of centriolar satellites, indicating disruption of the satellite compartment [36]. Consistent with this, GL2 control cells showed compact PCM1 signal around γ-tubulin-positive centrosomes, whereas depletion of Sld5 using two independent RNAi reagents caused marked PCM1 dispersion throughout the cytoplasm (Fig. 2A, D). We next asked whether Sld5 loss also disrupts additional centriolar satellite components. AZI1, which is required for centrosome duplication and whose pericentrosomal localization depends on PCM1 [26], was similarly dispersed in Sld5-depleted cells (Fig. 2B, D). CEP290, a centriolar satellite protein involved in centrosomal protein recruitment [23], also lost its compact centrosome-associated distribution following Sld5 depletion (Fig. 2C, D). Importantly, immunoblotting showed that the total cellular abundance of AZI1 and CEP290 was not substantially altered after Sld5 knockdown (Fig. 2E). Thus, Sld5 depletion does not simply reduce satellite protein expression, but instead disrupts the spatial organization of the centriolar satellite compartment. Because centriolar satellites depend on microtubule-based trafficking and the dynein–dynactin system for pericentrosomal localization, we next tested whether dispersal of AZI1 or CEP290 was sufficient to explain the spindle-pole defects observed in Sld5-deficient cells. Sld5 depletion caused a pronounced increase in multipolar spindle formation and γ-tubulin misorganization (Fig. 2F, G). By contrast, depletion of AZI1 or CEP290 produced substantially weaker spindle-pole defects and did not phenocopy the severity of the Sld5-deficient phenotype (Fig. 2F, G). Immunoblotting confirmed efficient depletion of Sld5, AZI1, and CEP290 under the corresponding RNAi conditions (Fig. 2H). These findings indicate that the spindle-pole abnormalities caused by Sld5 loss are not primarily driven by loss or dispersal of AZI1 and CEP290 alone. Rather, Sld5 appears to control a broader centrosome-satellite axis required for maintaining centriolar satellite integrity and mitotic spindle-pole organization. Centriolar satellite positioning depends on an intact microtubule network. To determine whether the satellite defects in Sld5-depleted cells were solely due to impaired microtubule organization, we examined satellite behavior after microtubule depolymerization and regrowth. Nocodazole treatment disrupted the microtubule network and caused PCM1 dispersion, confirming that satellite organization is strongly dependent on microtubule integrity (Supplementary Fig. 2A-C). After nocodazole washout, GL2 control cells efficiently reassembled microtubules and restored compact PCM1 organization near the centrosome. In contrast, Sld5-depleted cells showed microtubule regrowth but failed to restore normal PCM1 satellite organization (Supplementary Fig. 2A-C). These observations indicate that Sld5 is required for centriolar satellite reassembly even when microtubule nucleation is restored. Thus, the Sld5-dependent satellite defect is unlikely to reflect only a failure of microtubule regrowth. Instead, Sld5 may regulate satellite integrity through an additional mechanism, potentially involving centrosome-directed satellite trafficking through dynein–dynactin-dependent transport.

**Fig. 2.**
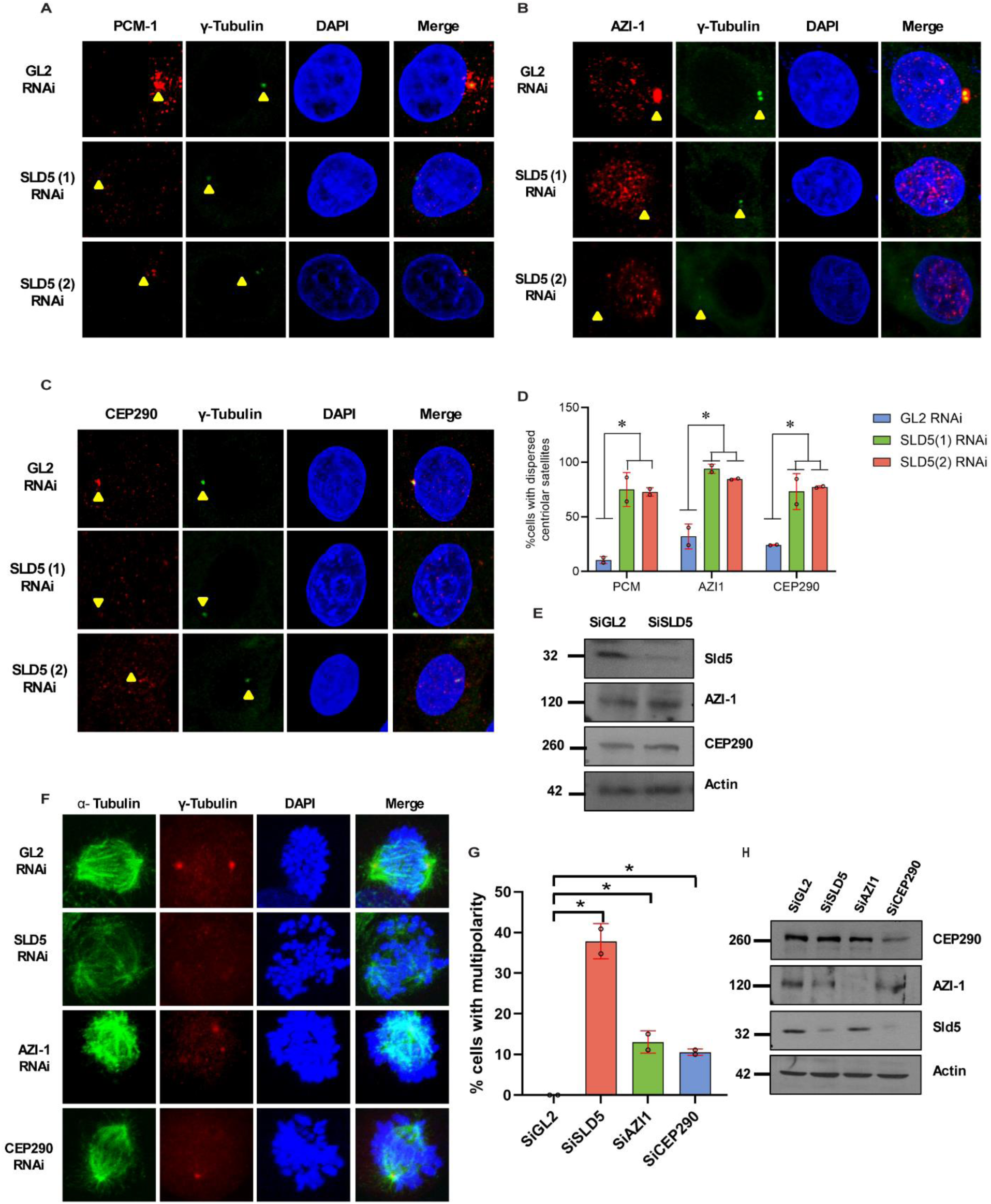
Sld5 depletion disperses centriolar satellite proteins and induces spindle-pole defects. **A-C.** Representative immunofluorescence images showing the localization of centriolar satellite proteins in GL2 control and Sld5-depleted cells. Cells were stained for PCM1 (A), AZI1 (B), or CEP290 (C) together with γ-tubulin to mark centrosomes and DAPI to mark DNA. Yellow arrowheads indicate γ-tubulin-positive centrosomal regions. In GL2 control cells, PCM1, AZI1, and CEP290 are enriched around centrosomes, whereas Sld5 depletion using two independent RNAi reagents causes dispersion of these centriolar satellite proteins. **D.** Quantification of cells showing dispersed PCM1, AZI1, or CEP290 signal after GL2 control RNAi or Sld5 RNAi. Sld5 depletion significantly increases the dispersion of all three centriolar satellite markers. Dots indicate individual measurements; error bars indicate variation among measurements; *P < 0.05. **E.** Immunoblot analysis of Sld5, AZI1, and CEP290 protein levels in GL2 control and Sld5-depleted cells. Actin was used as a loading control. Sld5 knockdown does not substantially reduce total AZI1 or CEP290 protein abundance, indicating that satellite disruption reflects altered localization rather than loss of protein expression. **F.** Representative mitotic cells stained for α-tubulin, γ-tubulin, and DAPI following GL2, Sld5, AZI1, or CEP290 RNAi. Sld5-depleted cells show prominent spindle-pole abnormalities and multipolarity, whereas AZI1- or CEP290-depleted cells show less severe spindle defects. **G.** Quantification of cells with multipolar spindles after GL2, Sld5, AZI1, or CEP290 RNAi. Sld5 depletion produces a significantly higher frequency of multipolarity than AZI1 or CEP290 depletion. Dots indicate individual measurements; error bars indicate variation among measurements; *P < 0.05. **H.** Immunoblot validation of CEP290, AZI1, and Sld5 depletion in the indicated RNAi conditions. Actin was used as a loading control.

### Sld5 couples centriolar satellite integrity to dynein–dynactin transport

Because centriolar satellites disperse away from centrosomes in Sld5-deficient cells, we next asked whether this defect reflects impaired dynein–dynactin-dependent trafficking. In cancer datasets, the expression of GINS4/SLD5 showed cancer-lineage-specific associations with DYNC1H1(DHC), which encodes the cytoplasmic dynein heavy chain. Analysis of SLD5 knockout-associated changes revealed variable relationships with DYNC1H1 across tumor types, including inverse associations in acute myeloid leukemia, hepatocellular carcinoma, sarcoma, and osteosarcoma, and positive associations in thyroid gland carcinoma and hematological malignancies (Fig. 3A and Supplementary Table S6). In parallel, analysis of SLD5-overexpressing tumor contexts showed positive associations between SLD5 and DYNC1H1 in several lineages, including esophageal, colorectal, bladder, ovarian, breast, and pancreatic carcinomas, whereas selected lineages showed weaker or inverse relationships (Fig. 3B and Supplementary Table S7). These data suggested that the GINS4/SLD5–DYNC1H1 axis may be regulated in a tumor-context-dependent manner. We then tested whether loss of Sld5 directly affects the dynein machinery in cells. Cytoplasmic dynein is a multi-subunit motor complex composed of a motor-domain-containing heavy chain, intermediate chains, light-intermediate chains, and light chains. Immunoblot analysis showed that depletion of Sld5 reduced the abundance of the dynein heavy chain, DHC/DYNC1H1, indicating that Sld5 deficiency compromises the core motor component of the complex (Fig. 3C). Consistent with this, transcript analysis showed significant downregulation of DHC mRNA after Sld5 depletion, whereas other dynein subunits showed variable degrees of regulation (Fig. 3D). These findings suggest that Sld5 loss preferentially affects DHC expression and may thereby destabilize dynein-complex function. We next examined the subcellular localization of dynein and dynactin components during mitosis. In GL2 control cells, DHC and LC1 were detected at γ-tubulin-positive spindle poles. In contrast, Sld5 depletion markedly reduced the spindle-pole localization of both DHC and LC1 (Fig. 3E,F,H,I). We further analyzed DCTN2/p50, also known as dynamitin, a dynactin subunit required for dynactin-complex assembly and cargo association [66]. Sld5-depleted cells showed reduced dynamitin (p50/DCTN2) signal at spindle poles, indicating impaired recruitment or retention of the dynactin machinery at centrosomal regions (Fig. 3G,J). Together, these data show that Sld5 is required for proper dynein–dynactin localization at spindle poles and support a model in which Sld5 deficiency disrupts centrosome-directed transport of satellite and centrosomal cargo.

**Fig. 3.**
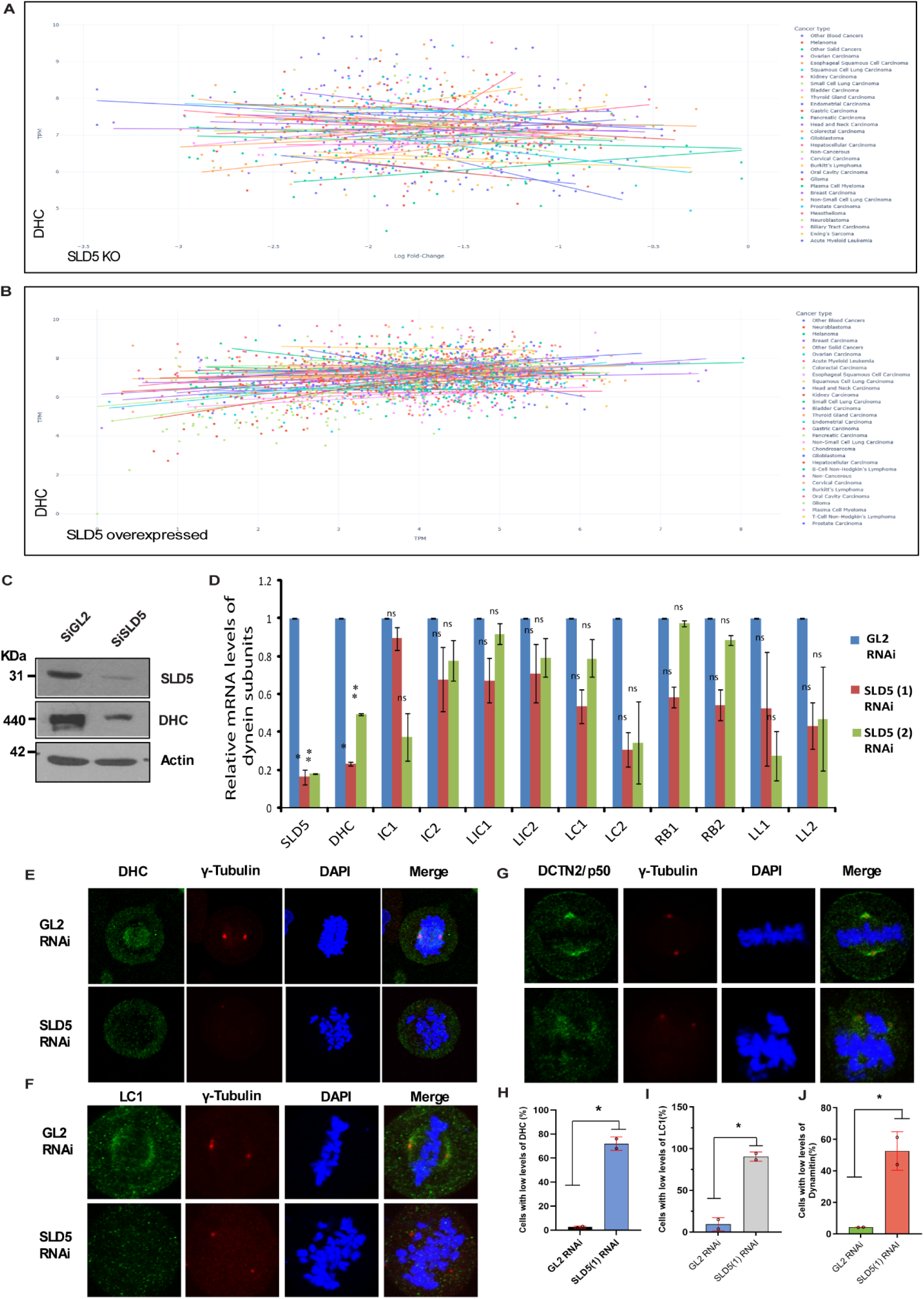
Sld5 depletion disrupts dynein–dynactin localization at spindle poles. **A.** Cancer-lineage-specific association between SLD5 knockout-associated log fold-change and DYNC1H1 expression. Each point represents a sample or cell line, and regression lines are shown for each cancer type. DYNC1H1 encodes the cytoplasmic dynein heavy chain. Summary correlation statistics are provided in Supplementary Table S6. **B.** Association between SLD5 expression in SLD5-overexpressing tumor contexts and DYNC1H1 expression across cancer types. Each point represents a sample or cell-line model, and regression lines are shown by cancer type. Summary correlation statistics are provided in Supplementary Table S7. **C.** Immunoblot analysis of Sld5 and dynein heavy chain in GL2 control and Sld5-depleted cells. Actin was used as a loading control. Sld5 depletion reduces DHC protein abundance. **D.** Relative mRNA expression of SLD5 and dynein-complex subunits after GL2 control RNAi or depletion of Sld5 using two independent RNAi reagents. Analyzed subunits include DHC, intermediate chains, light-intermediate chains, and light chains. Sld5 depletion significantly reduces DHC mRNA expression, whereas other dynein subunits show variable regulation. Bars show mean values with error bars; individual points indicate replicate measurements. ns, not significant; *P < 0.05; **P < 0.01. **E.** Representative immunofluorescence images showing DHC localization in GL2 control and Sld5-depleted mitotic cells. Cells were stained for DHC, γ-tubulin, and DAPI. Sld5 depletion reduces DHC signal at γ-tubulin-positive spindle poles. **F.** Representative immunofluorescence images showing LC1 localization in GL2 control and Sld5-depleted mitotic cells. Cells were stained for LC1, γ-tubulin, and DAPI. Sld5 depletion reduces LC1 accumulation at spindle poles. **G.** Representative immunofluorescence images showing DCTN2/p50, also known as dynamitin, in GL2 control and Sld5-depleted mitotic cells. Cells were stained for DCTN2/p50, γ-tubulin, and DAPI. Sld5 depletion reduces DCTN2/p50 localization at spindle poles. **H.** Quantification of cells with low DHC signal after GL2 control RNAi or Sld5 RNAi. Sld5 depletion significantly increases the proportion of cells with reduced DHC levels. *P < 0.05. **I.** Quantification of cells with low LC1 signal after GL2 control RNAi or Sld5 RNAi. Sld5 depletion significantly increases the proportion of cells with reduced LC1 levels. *P < 0.05. **J.** Quantification of cells with low dynamitin/DCTN2-p50 signal after GL2 control RNAi or Sld5 RNAi. Sld5 depletion significantly increases the proportion of cells with reduced dynamitin signal. *P < 0.05.

### Sld5-dependent spindle defects are not explained by DNA damage or checkpoint activation

Because Sld5 is a component of DNA replication machinery, we next determined whether the centrosomal and spindle-pole defects observed after Sld5 depletion were secondary to DNA damage or checkpoint activation. In both HeLa and U2OS cells, depletion of Sld5 did not induce detectable activation of DNA damage markers, including phospho-Chk1 and phospho-H2AX, whereas UV-treated positive-control cells showed robust marker induction (Supplementary Fig. S3A, B). Similarly, Sld5 overexpression did not trigger detectable DNA damage signaling compared with control transfections, whereas the UV-treated positive control confirmed assay responsiveness (Supplementary Fig. S3C). Analysis of phospho-histone H3 further indicated that Sld5 depletion did not cause marked accumulation of a mitotic checkpoint-associated signal under these conditions (Supplementary Fig. S3D). Protein-level tumor analysis also supported co-variation between GINS4/SLD5 and DYNC1H1 across tumor samples (Supplementary Fig. S3E). Collectively, these findings indicate that the dynein–dynactin and spindle-pole defects observed in Sld5-deficient cells are unlikely to result from a generalized DNA damage response.

### DHC depletion phenocopies Sld5 loss and disrupts centriolar satellite organization

Centriolar satellites are transported along microtubules towards centrosomes through the dynein–dynactin motor machinery, thereby enabling delivery of PCM1-associated protein cargo to the pericentrosomal region (Fig. 4A). Because Sld5 depletion caused both dynein–dynactin downregulation and centriolar satellite dispersion, we next asked whether loss of the dynein motor itself could account for the satellite and spindle-pole defects observed in Sld5-deficient cells. To test this, we compared control cells with cells depleted of either Sld5 or dynein heavy chain (DHC). Immunoblot analysis showed that Sld5 depletion reduced the abundance of several dynein-complex components, including DHC, dynein intermediate chains, light-intermediate chains, and light chains. Direct depletion of DHC produced a similar destabilization of dynein subunits, supporting the idea that loss of the DHC motor subunit compromises the integrity of the dynein complex (Fig. 4B). Consistent with the immunoblot data, immunofluorescence analysis showed that DHC localized prominently around γ-tubulin-positive spindle poles in GL2 control cells, whereas both Sld5 and DHC depletion caused a marked reduction in DHC signal at spindle poles (Fig. 4C, G). A similar phenotype was observed for LC1, a dynein light-chain component. LC1 was enriched near spindle poles in control cells but was strongly reduced after either Sld5 or DHC depletion (Fig. 4D, H). These data indicate that Sld5-dependent DHC loss impairs dynein-complex localization at centrosomal/spindle-pole regions. We next examined whether DHC depletion also reproduced the mitotic spindle defects caused by Sld5 loss. In control cells, α-tubulin formed a bipolar spindle with focused γ-tubulin-positive poles. In contrast, both Sld5- and DHC-depleted cells showed defective spindle architecture, reduced or asymmetric γ-tubulin signal, and increased spindle multipolarity (Fig. 4E, I). Thus, loss of DHC is sufficient to generate spindle-pole abnormalities resembling those observed after Sld5 depletion. Finally, we assessed whether DHC depletion also disrupted centriolar satellite organization. PCM1 was concentrated around centrosomes in GL2 control cells, whereas both Sld5 and DHC depletion caused extensive PCM1 dispersion throughout the cytoplasm (Fig. 4F, J). This phenocopy suggests that the centriolar satellite defects observed in Sld5-deficient cells arise, at least in major part, from impaired dynein-dependent transport. Supporting this interpretation, phase-contrast imaging showed similar cellular perturbation after Sld5 and DHC depletion (Supplementary Fig. S4A). In contrast, the Golgi marker giantin retained its compact perinuclear organization after Sld5 depletion, indicating that Sld5 loss does not globally disrupt all pericentrosomal or membrane-associated structures (Supplementary Fig. S4B). Together, these findings support a model in which Sld5 maintains centriolar satellite organization and spindle-pole integrity by preserving DHC-dependent dynein transport to the centrosome.

**Fig. 4.**
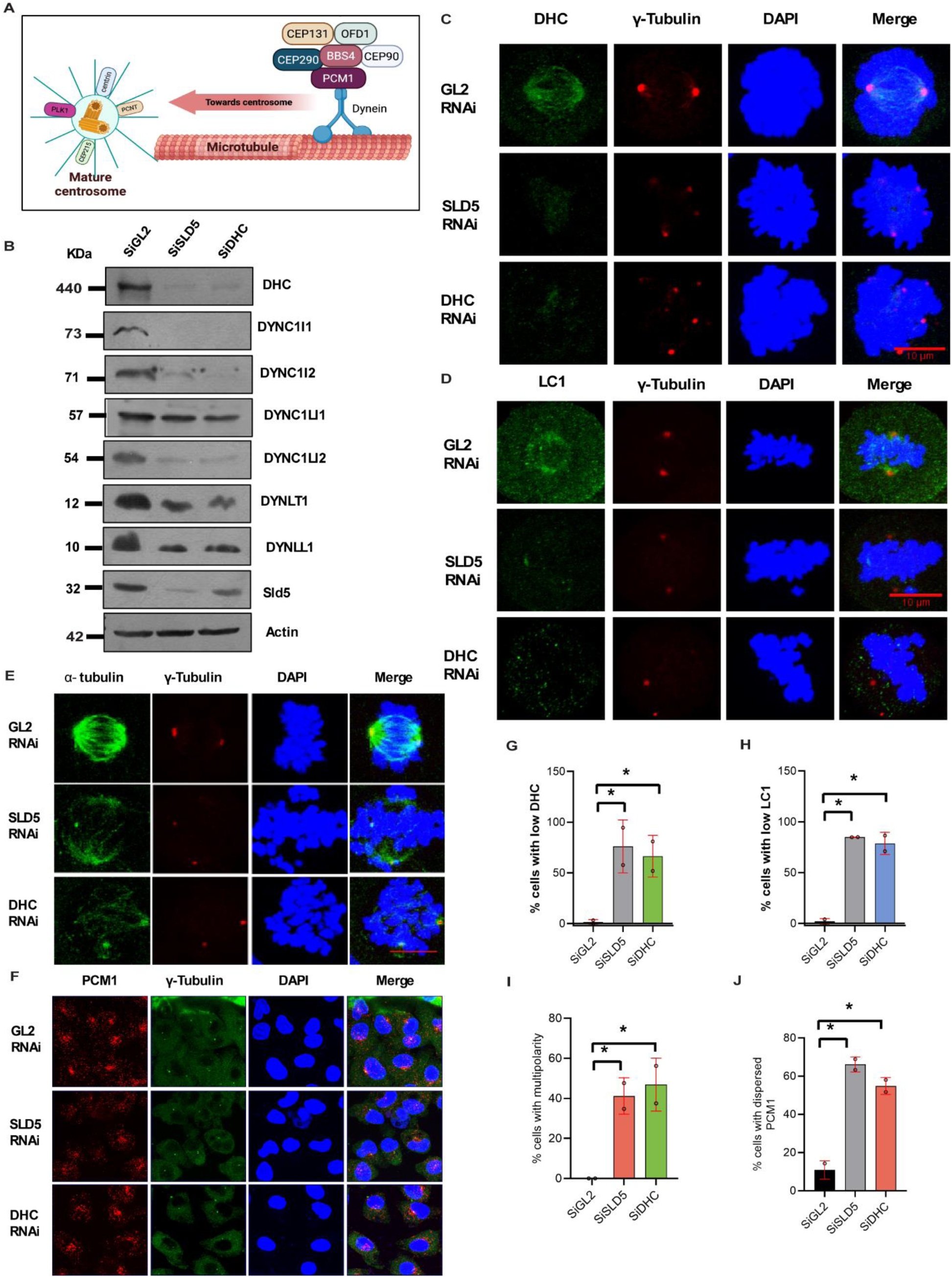
DHC depletion phenocopies Sld5 loss and causes centriolar satellite dispersion and spindle-pole defects. **A.** Schematic model showing dynein-dependent transport of centriolar satellite cargo towards the centrosome. PCM1-containing centriolar satellites recruit centrosomal proteins, including CEP131, OFD1, BBS4, and CEP290, and are transported along microtubules by dynein towards the mature centrosome. **B.** Immunoblot analysis of dynein-complex subunits in GL2 control, Sld5-depleted, and DHC-depleted cells. DHC, dynein intermediate chains, dynein light-intermediate chains, and dynein light chains are reduced after Sld5 depletion and are similarly affected after DHC depletion. Actin was used as a loading control. **C.** Representative immunofluorescence images showing DHC localization in GL2 control, Sld5-depleted, and DHC-depleted mitotic cells. Cells were stained for DHC, γ-tubulin, and DAPI. DHC signal is enriched at γ-tubulin-positive spindle poles in control cells but is reduced after Sld5 or DHC depletion. Scale bar, 10 μm. **D.** Representative immunofluorescence images showing LC1 localization in GL2 control, Sld5-depleted, and DHC-depleted mitotic cells. Cells were stained for LC1, γ-tubulin, and DAPI. LC1 localization at spindle poles is reduced after both Sld5 and DHC depletion. Scale bar, 10 μm. **E.** Representative immunofluorescence images of mitotic spindle organization after GL2 control, Sld5 or DHC RNAi. Cells were stained for α-tubulin, γ-tubulin, and DAPI. Control cells show bipolar spindle organization, whereas Sld5-and DHC-depleted cells show disrupted spindle architecture and abnormal γ-tubulin-positive spindle poles. **F.** Quantification of cells with low DHC signal after GL2 control, Sld5, or DHC RNAi. Both Sld5 and DHC depletion significantly increase the proportion of cells with reduced DHC signal. Dots indicate independent measurements; error bars indicate variation among measurements; *P < 0.05. **G.** Quantification of cells with low LC1 signal after GL2 control, Sld5, or DHC RNAi. Sld5 and DHC depletion significantly increase the proportion of cells with reduced LC1 signal. Dots indicate independent measurements; error bars indicate variation among measurements; *P < 0.05. **H.** Representative immunofluorescence images showing PCM1 localization after GL2 control, Sld5, or DHC RNAi. Cells were stained for PCM1, γ-tubulin, and DAPI. PCM1 is concentrated around centrosomal regions in control cells but becomes dispersed after Sld5 or DHC depletion. **I.** Quantification of cells with multipolar spindles after GL2 control, Sld5, or DHC RNAi. Sld5 and DHC depletion significantly increase spindle multipolarity. Dots indicate independent measurements; error bars indicate variation among measurements; *P < 0.05. **J.** Quantification of cells with dispersed PCM1 after GL2 control, Sld5, or DHC RNAi. DHC depletion causes PCM1 dispersion similar to that observed after Sld5 depletion. Dots indicate independent measurements; error bars indicate variation among measurements; *P < 0.05.

### Sld5-dependent DHC function is required for centrosome maturation

Centriolar satellites deliver centrosomal protein cargo along microtubules through dynein-dependent transport, thereby supporting centrosome growth and maturation before mitosis. Because Sld5 depletion caused both downregulation of dynein–dynactin and centriolar satellite dispersion, we next asked whether this defect impaired the recruitment of centrosome maturation factors. In GL2 control cells, PLK1, Aurora A, and CEP192 were efficiently concentrated at γ-tubulin-positive centrosomes during G2 phase. By contrast, depletion of either Sld5 or DHC markedly reduced centrosomal accumulation of these maturation-associated proteins (Fig. 5A-C). Quantification confirmed a significant reduction in the mean centrosomal intensity of PLK1, Aurora A, and CEP192 in both Sld5- and DHC-depleted cells compared with control cells (Fig. 5D-F). Thus, loss of Sld5 or DHC compromises the recruitment of key centrosome maturation factors during interphase. We then examined whether this maturation defect persisted during mitosis. In mitotic cells in control, PLK1, Aurora A, and CEP192 were enriched at γ-tubulin-positive spindle poles. In contrast, Sld5- and DHC-depleted cells showed reduced spindle-pole localization of PLK1, Aurora A, and CEP192, together with abnormal spindle-pole organization (Fig. 5G-I). Quantitative analysis showed a significant increase in cells with low mitotic PLK1, Aurora A, and CEP192 signal after either Sld5 or DHC depletion (Fig. 5J-L). These results indicate that Sld5-dependent maintenance of DHC is required for centrosome maturation both before and during mitosis. Because centrosome maturation depends on the delivery of protein cargo to centrosomes, these findings support a model in which Sld5 depletion reduces DHC-dependent transport, thereby impairing the movement of centriolar satellite cargo to centrosomes. Poorly matured centrosomes may then fail to withstand mitotic spindle forces, resulting in spindle-pole fragmentation and multipolarity.

**Fig. 5.**
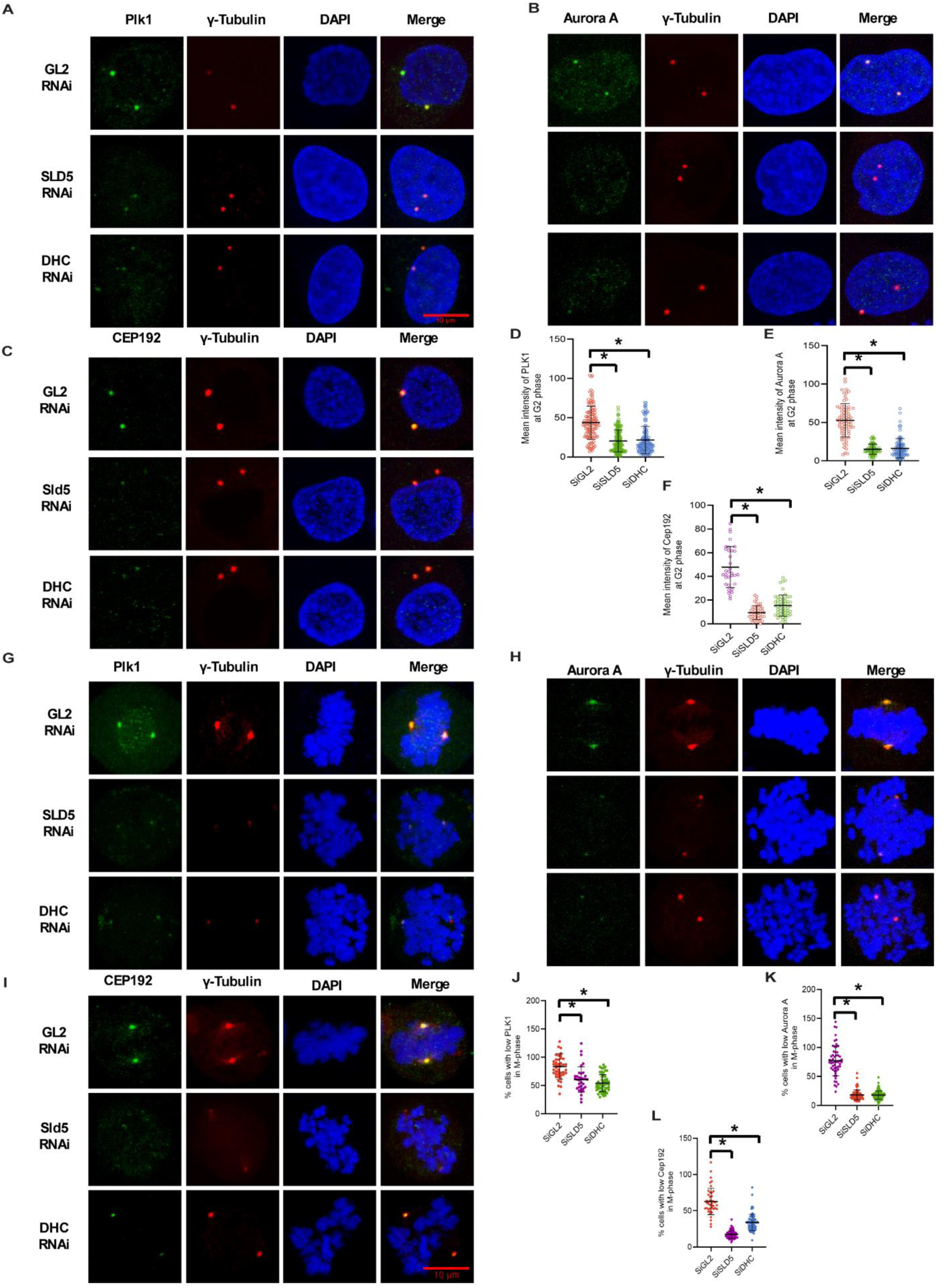
Sld5-dependent DHC function is required for centrosome maturation during interphase and mitosis. **A-C.** Representative immunofluorescence images showing centrosome maturation factors in G2-phase cells after GL2 control, Sld5, or DHC RNAi. Cells were stained for PLK1 (A), Aurora A (B), or CEP192 (C) together with γ-tubulin and DAPI. In control cells, PLK1, Aurora A, and CEP192 are enriched at γ-tubulin-positive centrosomes, whereas depletion of Sld5 or DHC reduces their centrosomal recruitment. **D-F.** Quantification of centrosomal signal intensity for PLK1 (D), Aurora A (E), and CEP192 (F) in G2-phase cells. Sld5 and DHC depletion significantly reduce the mean centrosomal intensity of all three maturation-associated proteins. Individual points represent quantified cells; bars indicate mean values with variation; *P < 0.05. **G-I.** Representative immunofluorescence images showing mitotic localization of PLK1 (G), Aurora A (H), and CEP192 (I) after GL2 control, Sld5, or DHC RNAi. Cells were stained with the indicated maturation marker, γ-tubulin, and DAPI. Control cells show enrichment of maturation factors at spindle poles, whereas Sld5- and DHC-depleted cells show reduced spindle-pole localization and abnormal spindle-pole organization. **J-L.** Quantification of mitotic cells with low PLK1 (J), Aurora A (K), or CEP192 (L) signal at spindle poles. Sld5 and DHC depletion significantly increase the proportion of cells with reduced mitotic centrosome maturation markers. Individual points represent quantified cells; bars indicate mean values with variation; *P < 0.05.

### Sld5 and DHC act within a shared centrosome–satellite pathway

To determine whether Sld5 and DHC function in the same pathway, we compared the effects of individual depletion of Sld5 or DHC with those of combined depletion of both proteins. Immunoblotting and RT–qPCR confirmed depletion of Sld5 and DHC at the protein and transcript levels, respectively (Supplementary Fig. S5A, B). Co-depletion of Sld5 and DHC did not produce an additive increase in centriolar satellite dispersion compared with individual depletion, as assessed by PCM1 localization (Supplementary Fig. S5C, G). Similarly, loss of DYNLT1 localization, reduced CEP215 recruitment, and spindle-pole multipolarity were not exacerbated by combined Sld5 and DHC depletion beyond the effects observed after single depletion (Supplementary Fig. S5D-J). These results indicate that Sld5 and DHC operate within a common pathway that controls centriolar satellite organization, dynein-dependent cargo transport, and centrosome maturation. Together with the phenocopy observed after DHC depletion, the data supports the conclusion that Sld5 loss disrupts centrosome integrity primarily by downregulating the DHC-dependent dynein transport machinery.

### Dynein motor activity is required for centriolar satellite organization and spindle-pole integrity

To determine whether the motor activity of dynein is required for centriolar satellite organization, we inhibited cytoplasmic dynein using ciliobrevin D, a small-molecule inhibitor that targets the ATPase activity of the dynein motor domain [67]. Cells were arrested in metaphase with MG132 and then treated with ciliobrevin D before analysis of spindle-pole architecture and centriolar satellite organization (Fig. 6A). In vehicle-treated cells, α-tubulin formed a compact bipolar spindle, and γ-tubulin localized to focused spindle poles. By contrast, ciliobrevin D treatment caused prominent spindle-pole defects, including disorganized microtubules and abnormal γ-tubulin-positive poles (Fig. 6B). Quantification confirmed a significant increase in cells with spindle-pole defects after dynein inhibition (Fig. 6D). These defects closely resembled the phenotypes observed after Sld5 or DHC depletion, indicating that dynein motor activity is required to maintain mitotic spindle-pole integrity. We next examined whether inhibition of dynein motor activity also affected centriolar satellites. In DMSO-treated cells, PCM1 was concentrated around the centrosomal region. Ciliobrevin D treatment caused marked dispersion of PCM1 throughout the cytoplasm, consistent with disruption of dynein-dependent centriolar satellite transport (Fig. 6C). Quantitative analysis showed a significant increase in cells with dispersed centriolar satellites following ciliobrevin D treatment (Fig. 6E). These data demonstrate that dynein motor activity is essential for centriolar satellite organization and that inhibition of dynein is sufficient to reproduce the satellite and spindle-pole defects observed in Sld5-deficient cells. Because Sld5 depletion reduced DHC expression and impaired centrosome-directed transport, we next integrated these observations into a working model. In Sld5-proficient cells, Sld5 supports dynein expression and dynein-dependent transport of PCM1-associated centriolar satellite cargo towards centrosomes, enabling centrosome maturation and bipolar spindle formation. In Sld5-deficient cells, DHC downregulation compromises dynein transport, leading to centriolar satellite dispersion, reduced delivery of centrosomal cargo, and formation of immature centrosomes that fail to maintain spindle-pole integrity during mitosis (Fig. 6F). To explore a potential upstream mechanism linking Sld5 to dynein expression, we examined the relationship between GINS4/SLD5 and POLR2A, which encodes the largest subunit of RNA polymerase II. Pan-cancer correlation analysis showed coordinated expression of GINS4 and POLR2A in multiple tumor lineages, with strong positive correlations in prostate carcinoma, B-lymphoblastic leukemia, other blood cancers, ovarian carcinoma, kidney carcinoma, and colorectal carcinoma (Fig. 6G and Table S8). Protein-level tumor analysis further supported variation in GINS4 abundance across tumor groups in the same expression framework (Fig. 6H). Structural modeling predicted a potential GINS4–POLR2A interface, and immunoprecipitation confirmed that RNA polymerase II was detected in Sld5 immunoprecipitation and not in the IgG control (Fig. 6I-K). Together, these observations suggest that Sld5 may interface with RNA polymerase II-associated transcriptional machinery, providing a possible mechanism by which Sld5 influences DHC expression and dynein-dependent centrosome regulation.

**Fig. 6.**
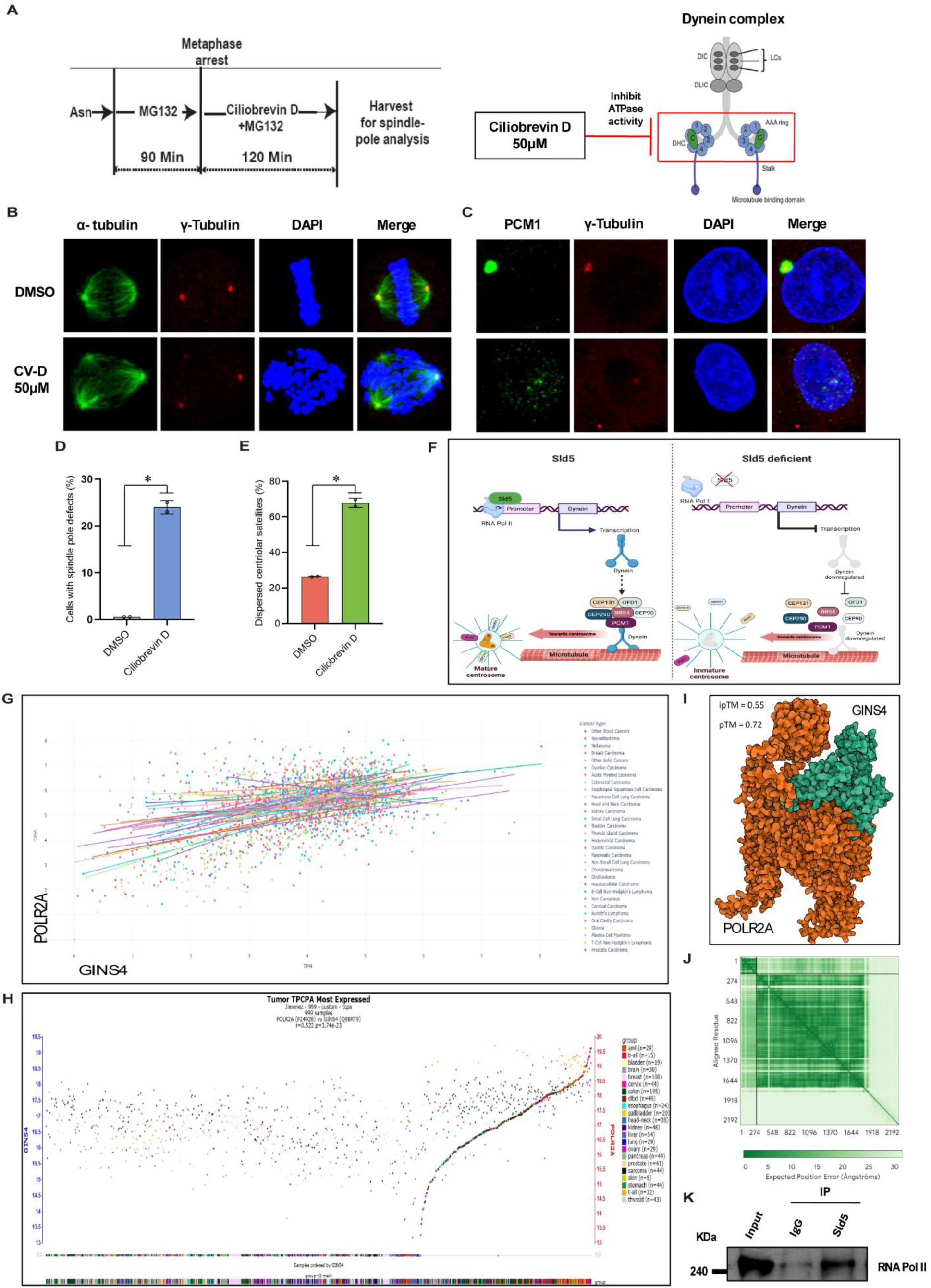
Dynein motor inhibition phenocopies Sld5 deficiency and disrupts centriolar satellite organization. **A.** Experimental scheme for dynein motor inhibition. Cells were arrested in metaphase with MG132 and subsequently treated with ciliobrevin D in the continued presence of MG132 before harvesting for spindle-pole analysis. The schematic on the right shows ciliobrevin D-mediated inhibition of dynein ATPase activity. **B.** Representative immunofluorescence images showing spindle organization in DMSO-treated and ciliobrevin D-treated cells. Cells were stained for α-tubulin, γ-tubulin, and DAPI. DMSO-treated cells show bipolar spindle organization, whereas ciliobrevin D-treated cells show spindle-pole defects and abnormal γ-tubulin-positive pole organization. **C.** Representative immunofluorescence images showing PCM1 localization in DMSO-treated and ciliobrevin D-treated cells. Cells were stained for PCM1, γ-tubulin, and DAPI. PCM1 is concentrated near the centrosomal region in control cells but becomes dispersed after ciliobrevin D treatment. **D.** Quantification of cells with spindle-pole defects after DMSO or ciliobrevin D treatment. Dynein inhibition significantly increases the frequency of spindle-pole defects. Bars show mean values with replicate measurements; *P < 0.05. **E.** Quantification of cells with dispersed centriolar satellites after DMSO or ciliobrevin D treatment. Ciliobrevin D significantly increases PCM1 dispersion. Bars show mean values with replicate measurements; *P < 0.05. **F.** Proposed model linking Sld5 to dynein-dependent centriolar satellite transport and centrosome maturation. In Sld5-proficient cells, Sld5 supports dynein expression and promotes dynein-mediated transport of PCM1-associated satellite cargo to centrosomes. In Sld5-deficient cells, reduced DHC expression impairs dynein transport, leading to centriolar satellite dispersion, defective centrosome maturation, and spindle-pole abnormalities. **G.** Pan-cancer correlation analysis of GINS4/SLD5 and POLR2A expression across cancer types. Each point represents a sample or cell line, and regression lines are shown for each cancer type. Summary correlation statistics are provided in Supplementary Table S8. **H.** Tumor proteomic analysis showing GINS4 abundance across tumor samples coloured by tumor group, supporting tumor-context-dependent variation in GINS4 protein expression. **I.** Structural prediction of a potential interaction between GINS4 and POLR2A. The predicted complex is shown with GINS4 and POLR2A indicated. Model confidence values are shown as ipTM and pTM scores. **J.** Predicted aligned error plot corresponding to the GINS4–POLR2A structural model. **K.** Co-immunoprecipitation analysis of Sld5 and RNA polymerase II. RNA polymerase II was detected in Sld5 immunoprecipitates but not in IgG control, supporting an association between Sld5 and RNA polymerase II-associated machinery.

### SLD5 dependency defines a therapeutically exploitable centrosome-dynein vulnerability

The preceding data establish that Sld5 depletion disrupts the dynein–dynactin machinery, disperses PCM1-positive centriolar satellites, impairs centrosome maturation, and produces spindle-pole fragmentation. Because these defects were phenocopied by DHC depletion and by pharmacological inhibition of dynein motor activity, we next asked whether SLD5 could represent a therapeutically relevant vulnerability in cancer. Loss-of-function profiling showed that SLD5 perturbation produced broadly negative log fold-change values across cancer models, indicating reduced cellular fitness after SLD5 loss (Fig. 7A). The distribution of log fold-change values was centred below zero and remained negative across multiple cancer lineages, although the magnitude of dependency varied by tumor type (Fig. 7B). These data suggest that many cancers require SLD5 for sustained proliferation, with a subset of lineages showing stronger sensitivity to SLD5 loss. To identify druggable pathways that may intersect with SLD5 dependency, we next analyzed predicted kinase associations with GINS4/SLD5. This analysis prioritized several mitotic, checkpoint, and replication-associated kinases, including BUB1, WEE1, CDC7, EIF2AK1, VRK2, NEK2, PLK4, TTK, and PLK1 (Table S9). Consistent with a therapeutically relevant mitotic network, survival-based forest plot analysis showed that several candidate kinase nodes were associated with altered patient outcomes, with a subset showing hazard ratios greater than 1, consistent with adverse prognostic impact (Fig. 7C). Co-occurrence analysis further showed significant co-occurrence among several kinase candidates, including GINS4 with EIF2AK1, WEE1 and PLK1, supporting the existence of a GINS4-associated kinase vulnerability module (Table S10). Together with the cell biological data, these results support a therapeutic model in which SLD5 maintains dynein-dependent centriolar satellite trafficking and centrosome maturation. In SLD5-proficient cells, PCM1-associated centriolar satellites are transported along microtubules by dynein to deliver centrosomal cargo, enabling centrosome maturation and bipolar spindle formation. In SLD5-deficient cells, DHC downregulation compromises dynein transport, leading to centriolar satellite dispersion, defective recruitment of centrosome maturation factors, and the formation of immature centrosomes. Upon mitotic entry, these centrosomes fail to withstand spindle forces, resulting in abnormal chromosome attachments, spindle-pole fragmentation, and multipolar spindle formation (Fig. 7D). These findings nominate the SLD5–dynein–centrosome axis as a cancer vulnerability and suggest that SLD5-dependent tumors may be therapeutically sensitized by targeting cooperating mitotic and checkpoint kinases, including WEE1, CDC7, PLK1, BUB1, and related centrosome-associated regulators.

**Fig. 7.**
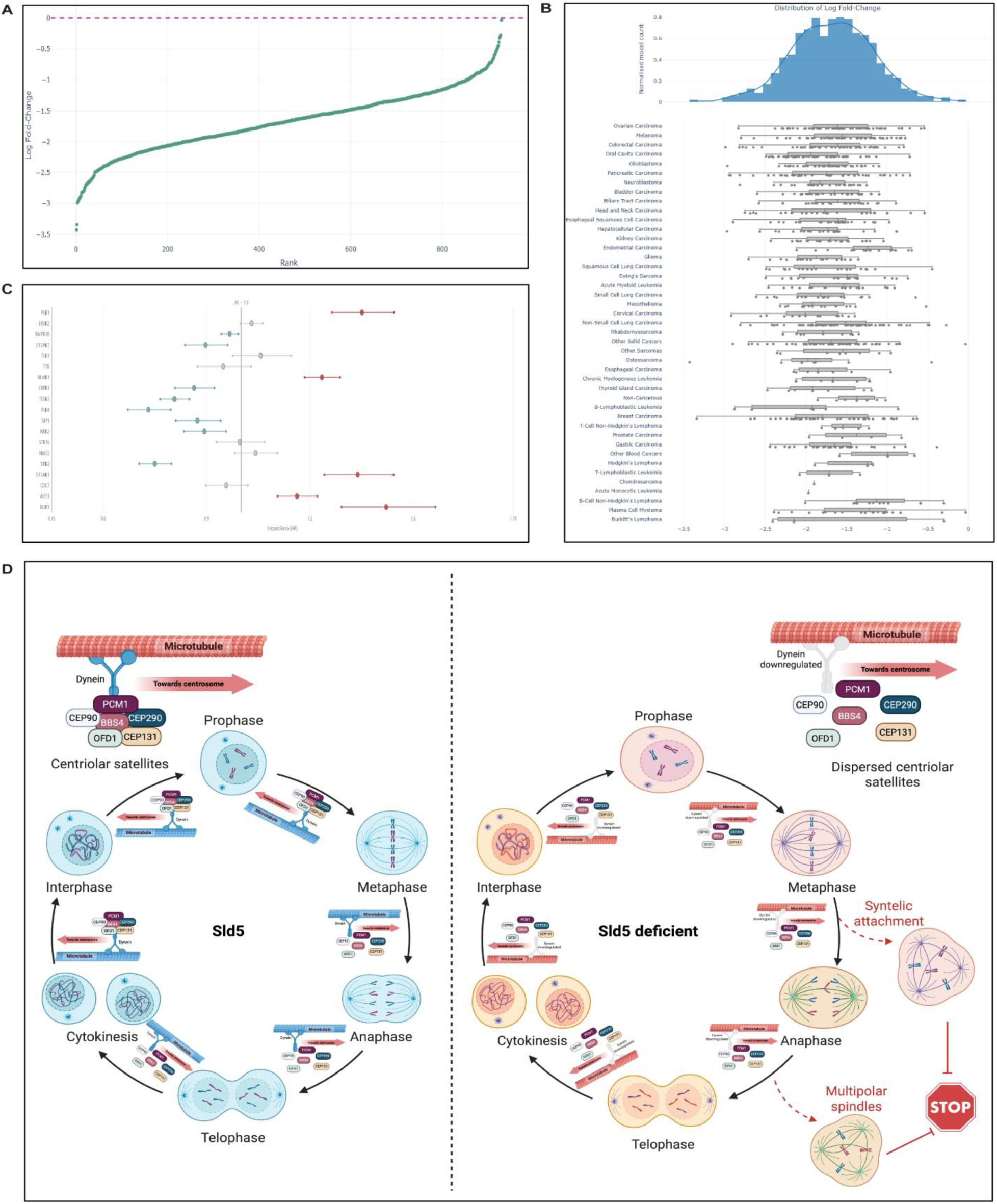
Therapeutic vulnerability framework for the SLD5-dynein-centrosome axis. **A.** Rank-ordered loss-of-function profile showing SLD5-associated log fold-change values across cancer models. The dashed line indicates no change. Negative log fold-change values indicate reduced fitness after SLD5 perturbation, supporting SLD5 as a broadly required cancer dependency. **B.** Distribution of SLD5 loss-of-function log fold-change values across cancer types. The upper histogram shows the global distribution of log fold-change values, and the lower panel shows cancer-lineage-specific distributions. SLD5 loss produces negative fitness effects across multiple tumor lineages, with lineage-dependent variation in the strength of dependency. **C.** Forest plot showing hazard ratios for candidate kinase genes associated with the GINS4/SLD5 therapeutic network. The vertical line denotes HR = 1. Points indicate estimated hazard ratios, and horizontal bars indicate confidence intervals. Candidate kinase nodes with HR > 1 are associated with increased risk, whereas nodes with HR < 1 are associated with reduced risk. These results identify mitotic, checkpoint, and replication-associated kinases as clinically relevant components of the SLD5-associated vulnerability network. **D.** Proposed therapeutic model. In SLD5-proficient cells, dynein-dependent transport moves PCM1-containing centriolar satellites and associated centrosomal cargo towards centrosomes, supporting centrosome maturation, bipolar spindle assembly, and normal mitotic progression. In SLD5-deficient cells, DHC downregulation impairs dynein-dependent transport, leading to centriolar satellite dispersion, defective centrosome maturation, and impaired spindle-pole integrity. These defects promote abnormal chromosome attachments, spindle-pole fragmentation and multipolar spindle formation, providing a therapeutic opportunity to target the SLD5–dynein–centrosome axis and cooperating mitotic kinase pathways.

## Discussion

This study identifies SLD5/GINS4 as a cancer-associated replication factor with an unanticipated role in centrosome biology. By integrating pan-cancer transcriptomic and proteomic analyses with mechanistic cell biology experiments, we show that SLD5 is recurrently upregulated in human cancers, supports dynein-dynactin-dependent centriolar satellite trafficking, and defines a therapeutically relevant centrosome-mitotic vulnerability. These findings extend the established role of SLD5 as a component of the GINS replication complex and place it within a broader cellular program that couples DNA replication, dynein-mediated cargo transport, centrosome maturation, and spindle-pole integrity.

GINS4, also known as SLD5, is one of the four subunits of the GINS complex, together with PSF1, PSF2, and PSF3. The GINS complex is an essential component of the eukaryotic replisome and participates in the initiation and elongation phases of DNA replication as part of the Cdc45–MCM–GINS helicase complex [27–30]. Because malignant cells frequently exhibit increased proliferative demand and replication stress, genes involved in replication licensing and fork progression are often transcriptionally activated in cancer. In this context, our pan-cancer analyses show that GINS4 is significantly elevated across multiple tumor types at the mRNA level and is also increased at the protein level in tumor proteomic and immunohistochemical datasets. The high Stouffer’s Z-score and positive cancer-specific enrichment signals further support the view that GINS4 upregulation is not restricted to a single tumor lineage but represents a recurrent pan-cancer event. These observations are consistent with previous reports implicating GINS4 in cancer progression. Pan-cancer analyses have reported GINS4 overexpression across many human malignancies and linked its abundance to prognosis, immune infiltration, and tumor-associated pathways [34, 35]. Tumor-specific studies have also associated increased GINS4 expression with poor clinical outcome and malignant progression in hepatocellular carcinoma, colorectal cancer, glioma, gastric cancer, and pancreatic cancer [31–33, 68, 69]. Our data extends these findings in two important ways. First, we provide a multi-layered expression framework combining mRNA, protein, immunohistochemistry, and integrated gene-ranking analyses. Second, we move beyond interpreting GINS4 as a mere proliferation marker by uncovering a centrosome–dynein mechanism that may explain how SLD5 supports cancer-cell fitness during mitotic progression. The pathway-level analyses further support this interpretation. GINS4-high tumor states were enriched for DNA replication, chromosome segregation, mitotic cell-cycle regulation, cytokinesis, sister chromatid cohesion, and spindle-associated programs. This profile is biologically coherent with the known replisome function of SLD5 but also suggests that GINS4-high tumors activate a broader replication–mitosis axis. Importantly, GINS4 expression also showed cancer-context-dependent associations with immune infiltration. This does not necessarily mean that GINS4 directly regulates immune recruitment; rather, it suggests that GINS4-high tumor states may coexist with distinct tumor microenvironment architectures. Because proliferation, aneuploidy, replication stress, and immune context are increasingly recognized as interconnected tumor features, these findings position GINS4 as a candidate biomarker of aggressive, replication-driven cancer states.

Although SLD5 is classically studied as a DNA replication protein, previous work from Kaur et al. showed that SLD5 localizes to centrosomes and helps preserve centriolar satellites, thereby enabling centrosomes to resist congression forces during mitosis [36]. Our work builds directly on this observation and defines the underlying mechanism. We show that SLD5 depletion disperses PCM1, AZI1, and CEP290, three important centriolar satellite-associated proteins, without markedly reducing total AZI1 or CEP290 protein abundance. Thus, SLD5 loss does not simply eliminate satellite proteins; rather, it disrupts their spatial organization. Centriolar satellites are dynamic, PCM1-containing pericentrosomal assemblies that act as trafficking hubs and protein reservoirs for centrosomes and cilia. PCM1 has been described as a core scaffold of the satellite compartment, and PCM1-containing satellites contribute to the microtubule- and dynactin-dependent recruitment of proteins to the centrosome [12, 15, 70]. Our nocodazole washout experiments show that control cells can reassemble centriolar satellites after microtubule regrowth, whereas SLD5-depleted cells fail to restore normal PCM1 organization despite recovery of microtubule nucleation. This result is important because it separates the SLD5 phenotype from a simple defect in microtubule regrowth. Instead, it suggests that SLD5 is required for the transport or retention machinery that repositions satellites around centrosomes. The central mechanistic finding of this study is that SLD5 preserves centriolar satellite organization by maintaining the dynein–dynactin transport system. Cytoplasmic dynein is the major minus-end-directed microtubule motor, and dynein–dynactin complexes are required for intracellular cargo transport, spindle organization, centrosome function, and spindle-pole focusing [22, 71–73]. We found that SLD5 depletion reduces DHC/DYNC1H1 abundance at both the mRNA and protein levels and destabilizes additional dynein-complex subunits. Consistent with these biochemical defects, SLD5-depleted cells show reduced spindle-pole localization of DHC, LC1, and DCTN2/p50/dynamitin. Because DCTN2/p50 is a dynactin subunit involved in dynein–dynactin organization and cargo association, its loss from spindle-pole regions further supports defective centrosome-directed transport. Several lines of evidence indicate that DHC downregulation is not merely associated with the SLD5 phenotype but functionally contributes to it. First, direct depletion of DHC phenocopied SLD5 depletion, causing destabilization of the dynein complex, reduced DHC and LC1 localization, PCM1 dispersion, spindle-pole defects, and multipolarity. Second, combined SLD5 and DHC depletion did not markedly exacerbate the defects seen after individual depletion, supporting the conclusion that both proteins act within a shared pathway. Third, pharmacological inhibition of dynein motor activity with ciliobrevin D caused PCM1 dispersion and spindle-pole defects similar to those observed after SLD5 or DHC depletion. Ciliobrevins were originally described as small-molecule inhibitors of cytoplasmic dynein ATPase activity and are widely used to interrogate dynein motor function [74, 75]. Together, these findings strongly support a model in which SLD5 acts upstream of DHC-dependent dynein transport to maintain centriolar satellite positioning and spindle-pole integrity. The consequences of disrupting this pathway become apparent during centrosome maturation. Centrosome maturation requires recruitment of pericentriolar material and mitotic regulators, including CEP192, Aurora A, and PLK1. CEP192 organizes an Aurora A–PLK1 signaling cascade that is essential for centrosome maturation and bipolar spindle assembly, while PLK1-mediated phosphorylation events contribute to pericentriolar material expansion and spindle-pole organization [4, 6]. In our study, SLD5 or DHC depletion reduced centrosomal recruitment of PLK1, Aurora A and CEP192 during interphase and mitosis. This suggests that SLD5-dependent dynein transport is required to deliver or maintain centrosome maturation cargo at γ-tubulin-positive centrosomes. When this transport axis fails, centrosomes enter mitosis in an immature or structurally weakened state, making them vulnerable to fragmentation under spindle forces and resulting in multipolar spindle formation. A key strength of the present study is that the spindle-pole phenotype is not simply attributed to canonical DNA damage signaling. Because SLD5 is a replisome component, one possible explanation was that SLD5 depletion causes replication failure, DNA damage, checkpoint activation, and secondary mitotic defects. However, phospho-Chk1 and phospho-H2AX were not detectably induced after SLD5 depletion or overexpression under the tested conditions, whereas positive controls responded as expected. Thus, although SLD5 remains an essential replication protein, the centrosome and dynein defects observed here appear to represent a separable, non-canonical function of SLD5 rather than a downstream consequence of overt DNA damage. This distinction is important because it expands the biological role of SLD5 from replication fork function to the spatial coordination of centrosome-associated trafficking. The POLR2A data provides a potential upstream explanation for how SLD5 controls DHC abundance. Pan-cancer correlation, structural modeling, and co-immunoprecipitation suggest that SLD5 may interface with RNA polymerase II-associated transcriptional machinery. This observation is still hypothesis-generating and should not yet be interpreted as a fully defined transcriptional mechanism. However, it provides a plausible route by which SLD5 could influence DYNC1H1 transcription and thereby maintain the dynein–dynactin complex. Future studies should determine whether SLD5 directly regulates DYNC1H1 promoter occupancy, RNA polymerase II recruitment, transcription elongation, or transcript stability. Rescue experiments restoring DHC in SLD5-depleted cells would also be important to determine the extent to which DHC loss is sufficient to account for the entire SLD5 phenotype.

The therapeutic implication of this study is that SLD5/GINS4 may mark a cancer-cell state that depends on coordinated replication, dynein transport, and centrosome maturation. Loss-of-function profiling showed broadly negative SLD5-associated log-fold changes across cancer models, indicating that many cancer cells lose fitness when SLD5 is perturbed. This is consistent with large-scale cancer dependency projects showing that genome-wide CRISPR and RNAi screens can identify lineage-specific and pan-cancer vulnerabilities [58, 59]. In our model, SLD5 dependency is not explained solely by its role in DNA replication. Instead, SLD5 loss also compromises dynein-dependent satellite trafficking, reduces centrosome maturation, and produces mitotic failure. This dual replication–mitosis function may explain why SLD5 depletion is detrimental to proliferating cancer cells. Targeting SLD5 directly may be challenging, as the GINS complex is required for normal DNA replication. However, many successful cancer strategies exploit differential dependence rather than absolute tumor specificity. Tumors with high GINS4 expression, elevated replication pressure, centrosome abnormalities, or dynein–centrosome adaptation may have a narrower tolerance for additional perturbation of mitotic or checkpoint pathways. Our kinase-association and co-occurrence analyses prioritize several candidate therapeutic nodes, including WEE1, CDC7, PLK1, BUB1, TTK, PLK4, NEK2, and related mitotic regulators. These candidates are biologically consistent with the SLD5 phenotype. WEE1 and CDC7 regulate checkpoint and replication-stress responses; PLK1, PLK4, Aurora A-associated pathways, and CEP192-dependent signaling regulate centrosome maturation, spindle assembly, and mitotic progression; BUB1 and TTK participate in spindle checkpoint control. Thus, SLD5-high or SLD5-dependent tumors may be vulnerable to combining targeting of replication stress and centrosome/mitotic control pathways. The centrosome component of this vulnerability is especially important. Centrosome abnormalities are common in cancer and can promote chromosomal instability, but they also create therapeutic liabilities. Cancer cells with centrosome amplification often survive by clustering centrosomes into pseudo-bipolar spindles; disruption of this adaptation can force multipolar division and cell death [7, 8]. Our data suggest a related but distinct vulnerability: SLD5 loss impairs centrosome maturation and dynein-dependent satellite delivery, producing immature or unstable centrosomes that fragment during mitosis. Therefore, SLD5-dependent tumors may be sensitive not only to replication-stress drugs but also to agents that interfere with centrosome maturation, spindle-pole maintenance, or centrosome-clustering mechanisms. At the same time, this therapeutic concept should be developed cautiously. Dynein itself is a broadly essential motor, and systemic dynein inhibition is unlikely to be tolerated as a conventional anticancer strategy. The value of the SLD5–DHC axis may therefore lie less in direct dynein inhibition and more in patient stratification and rational combinations. For example, GINS4-high tumors could be tested for sensitivity to WEE1, CDC7, PLK1, TTK, or PLK4 inhibitors; SLD5 perturbation could be combined with mitotic kinase inhibition to identify synthetic lethal interactions; and DYNC1H1 expression, PCM1 dispersion, or centrosome maturation markers could be evaluated as pharmacodynamic readouts. Such studies would determine whether SLD5 expression identifies tumors that are especially dependent on the replication–centrosome axis.

Several limitations should be considered. The pan-cancer analyses identify associations between GINS4, clinical outcome, immune context, and kinase networks, but these correlations do not establish causality. The mechanistic experiments strongly support an SLD5–DHC–centriolar satellite pathway, but broader validation across multiple cancer lineages, three-dimensional cultures, and in vivo models will be needed. The POLR2A interaction also requires deeper mechanistic testing, including chromatin-based assays and transcriptional rescue experiments. Finally, because SLD5 is a core replication-associated protein, therapeutic strategies must define a tumor-selective window, either through lineage-specific dependency, combination therapy, or biomarker-guided patient selection.

In summary, our study reframes SLD5/GINS4 as more than a replisome subunit. We propose that SLD5 supports cancer-cell proliferation through an integrated axis linking replication competence, RNA polymerase II-associated regulation of DHC expression, dynein–dynactin-dependent centriolar satellite trafficking, centrosome maturation, and mitotic spindle-pole integrity. In SLD5-proficient cells, dynein transports PCM1-associated satellite cargo to centrosomes, enabling recruitment of PLK1, Aurora A, CEP192, and CEP215 and supporting bipolar mitosis. In SLD5-deficient cells, DHC downregulation disrupts this transport pathway, disperses centriolar satellites, impairs centrosome maturation, and generates multipolar spindle defects. This mechanism provides a biological explanation for SLD5 dependency in cancer and nominates the SLD5–dynein–centrosome axis as a candidate therapeutic vulnerability in replication-driven tumors.

## Acknowledgments

We thank Dr Sandeep Saxena, Manpreet Kaur, Ritu Shekhar, Gargi Roy, Monika Singh, Antara Mondal, and Sunder Bisht for assisting in various parts of this paper. We would like to thank National Institute of Immunology, Government of India to support this work. I would also like to thank the Department of Biotechnology for providing fellowship. We would like to thank https://BioRender.com for graphical abstract (Created in https://BioRender.com 2026) https://BioRender.com/oh80r7t).

## Funding

No disclosure

## Authors’ Contribution

Vipin Kumar conceived the project, conducted all the experiments, analyzed the data, and written the manuscript. Vivek Singh did all the bioinformatics and clinical relevance of SLD5 and wrote the manuscript. Raksha Devi, Praveen Kumar and Tanushree Ghosh have assisted at various stages of this manuscript.

## Declarations

### Ethics approval and consent to participate

Not applicable.

### Consent for publication

All authors have read and approved the final version of this manuscript.

### Competing interests

The authors declare no competing interests.

### Supplementary information

Supplementary data includes five supplementary figures and ten supplementary tables.

## Abbreviations

PCM: Pericentriolar matrix
CS: centriolar satellite
MT: microtubule
MTBD: microtubule binding domain
𝞬TuRC: 𝞬-tubulin ring complex
PLK: polo like-kinase
CEP: centrosomal protein
MTOC: microtubule organizing center
CDK: cyclin dependent kinase
PCM1: pericentrin material 1
DHC: dynein heavy chain (DYNC1H1)
IC1: Intermediate chain 1 (DYNC1I1)
IC2: Intermediate chain 2 (DYNC1I2)
LIC1: light intermediate chain1 (DYNC1LI1)
LIC2: light intermediate chain 2 (DYNC1LI2)
LC 1: light chain 1 (DYNLT1)
LC 2: light chain 2 (DYNLT3)
RB1: DYNLRB1(Light chain roadblock)
RB2: DYNLRB2(Light chain roadblock)
LC8-1: DYNLL1 (Light chain LC8-1)
LC8-2: DYNLL2 (Light chain LC8-2)
p150Glued: DCTN 1 (p150)
p50: DCTN2 (Dynamitin)

## Supplementary figure legends

**Supplementary Fig. S1.**
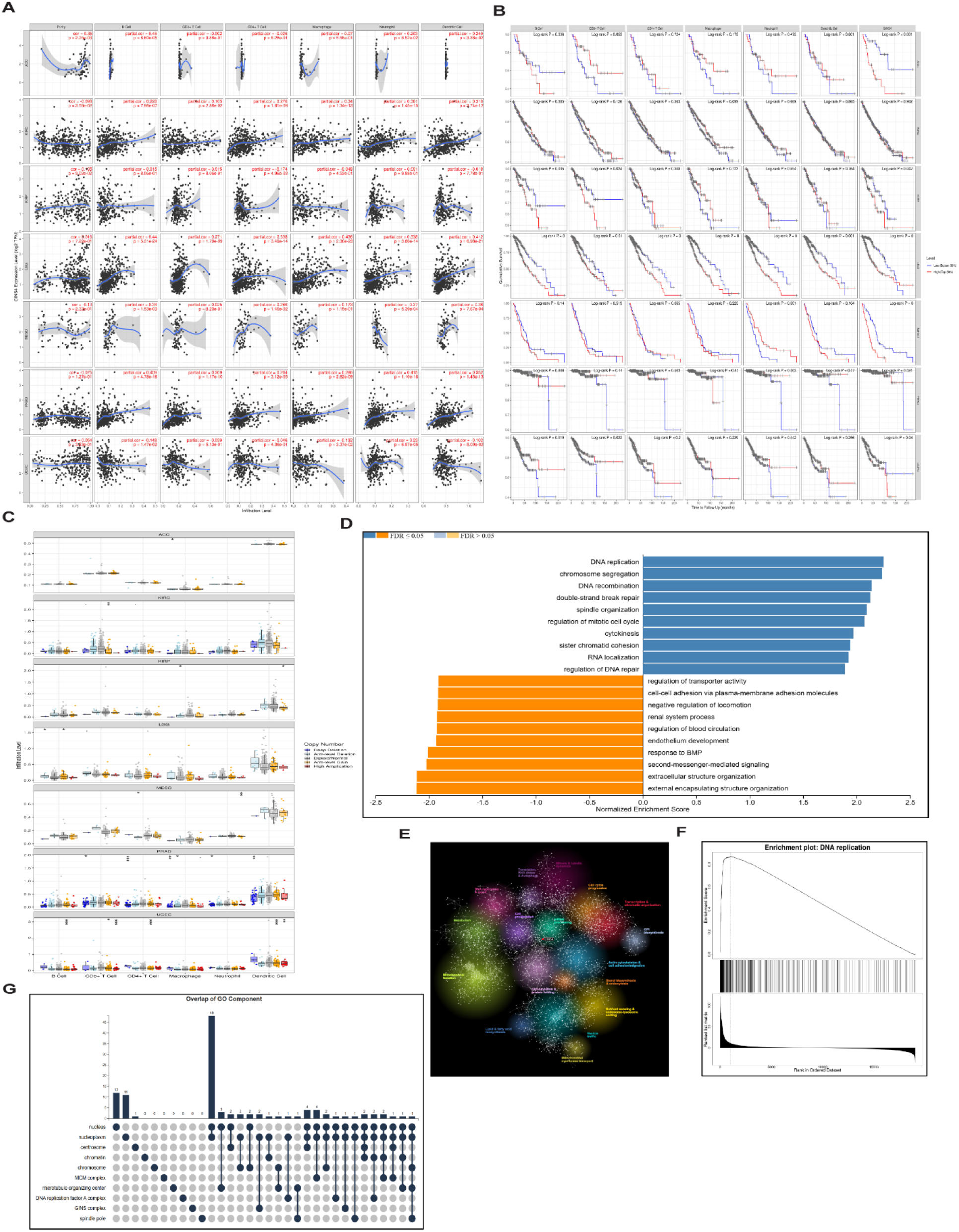
Immune-context and pathway analyses of GINS4-associated tumor states. **A.** Correlation analysis between GINS4 expression and tumor purity or immune-cell infiltration across selected cancer types. Immune compartments include B cells, CD8⁺ T cells, CD4⁺ T cells, macrophages, neutrophils, and dendritic cells. Correlation coefficients and P values are shown within each panel. **B.** Kaplan–Meier survival analyses stratified by high versus low GINS4 expression or immune-infiltration estimates across selected tumor contexts. Log-rank P values are shown in each panel. **C.** Association between GINS4 copy-number status and immune infiltration levels across selected cancer types. Copy-number categories include deep deletion, arm-level deletion, diploid/normal, arm-level gain, and high amplification. Box plots show the distribution of immune-infiltration estimates within each copy-number group. **D.** Gene set enrichment analysis of GINS4-associated transcriptional programs. Bars indicate normalized enrichment scores. Positively enriched pathways include DNA replication, chromosome segregation, DNA recombination, double-strand break repair, spindle organization, mitotic cell-cycle regulation, cytokinesis, and DNA repair regulation. Color denotes FDR status as indicated in the plot. **E.** Enrichment-map visualization of functionally related gene sets associated with GINS4-linked transcriptional programs. Gene-set clusters highlight biological processes associated with replication, mitosis, chromosomes, and DNA repair. **F.** Running enrichment plot for the DNA replication gene set, showing enrichment of DNA replication genes among the GINS4-associated ranked gene list. **G.** UpSet plot showing overlap among Gene Ontology cellular-component annotations for GINS4-associated genes. Enriched components include the nucleus, nucleoplasm, centrosome, chromatin, chromosome, MCM complex, microtubule-organizing center, DNA replication factor A complex, GINS complex, and spindle pole.

**Supplementary Fig. S2.**
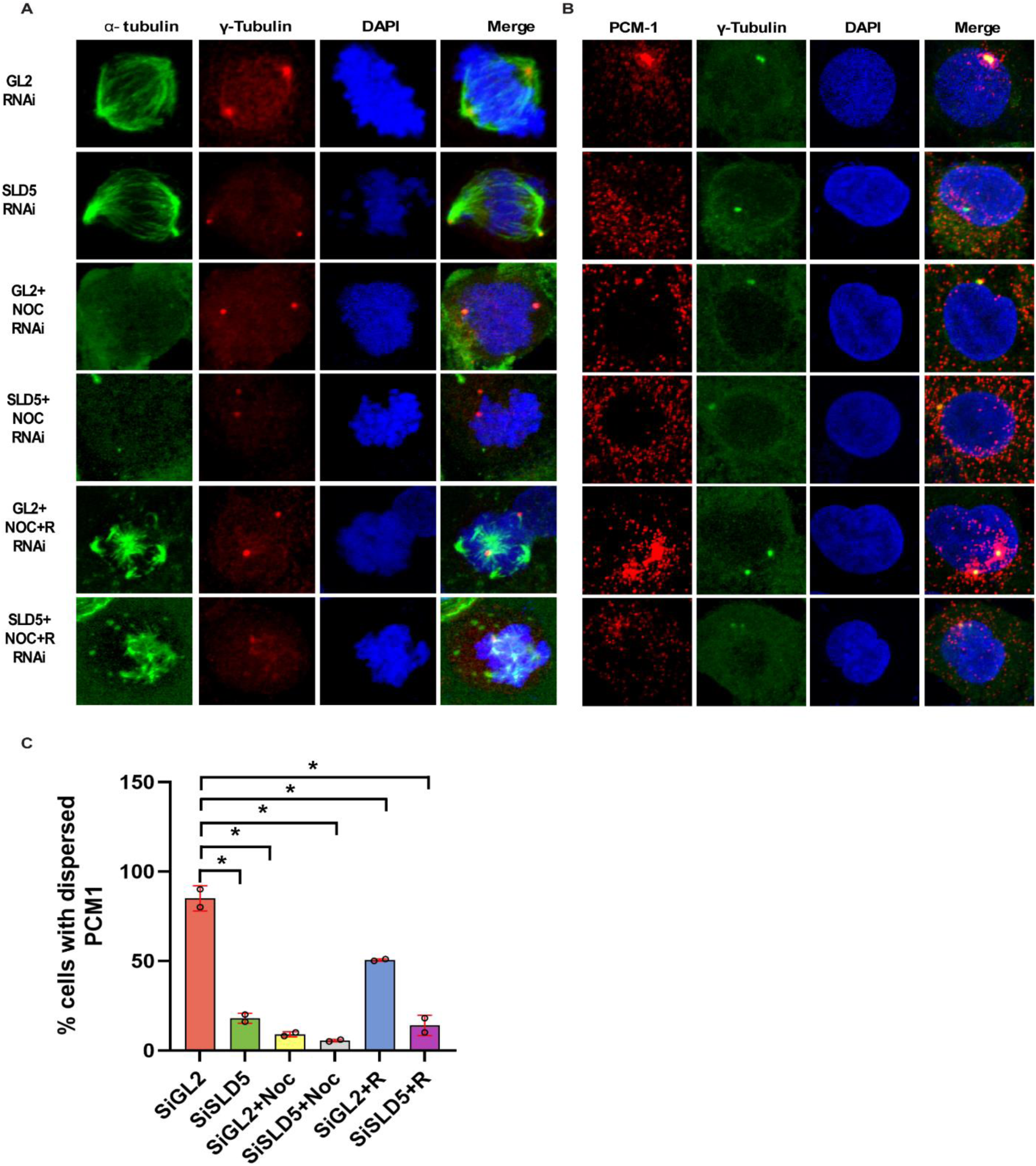
Sld5 depletion impairs centriolar satellite reassembly after microtubule regrowth. **A.** Representative immunofluorescence images showing α-tubulin, γ-tubulin, and DAPI staining in GL2 control and Sld5-depleted cells under basal conditions, after nocodazole treatment, and after nocodazole washout/recovery. Nocodazole disrupts the microtubule network, whereas recovery allows microtubule regrowth. **B.** Representative immunofluorescence images showing PCM1, γ-tubulin, and DAPI staining under the same conditions as in a. PCM1 is organized around centrosomes in GL2 control cells, becomes dispersed after nocodazole-induced microtubule depolymerization, and reassembles after microtubule recovery. In Sld5-depleted cells, PCM1 remains dispersed despite microtubule regrowth. **C.** Quantification of cells with dispersed PCM1 signal in GL2 control and Sld5-depleted cells under basal, nocodazole-treated, and recovery conditions. NOC denotes nocodazole treatment; R denotes recovery after nocodazole washout. Dots indicate individual measurements; error bars indicate variation among measurements; *P < 0.05.

**Supplementary Fig. S3.**
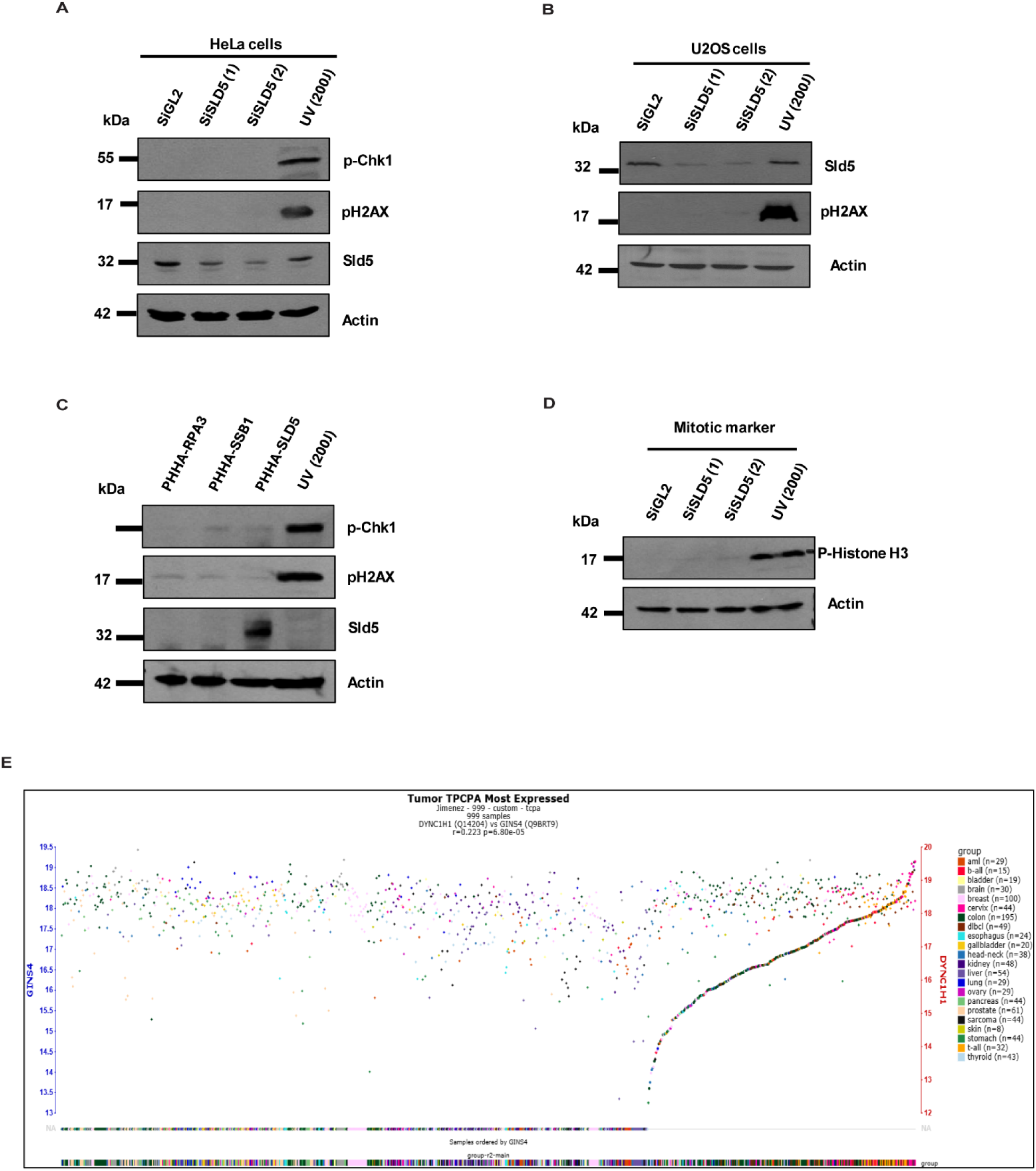
Sld5 depletion or overexpression does not induce detectable DNA damage signaling. **A.** Immunoblot analysis of phospho-Chk1, phospho-H2AX, Sld5, and actin in HeLa cells after GL2 control RNAi or Sld5 depletion using two independent RNAi reagents. UV-treated cells were used as a positive control for DNA damage induction. Sld5 depletion does not induce detectable phospho-Chk1 or phospho-H2AX accumulation. **B.** Immunoblot analysis of Sld5, phospho-H2AX, and actin in U2OS cells after GL2 control RNAi or Sld5 depletion. UV-treated cells were used as a positive control. Sld5 depletion does not induce detectable phospho-H2AX accumulation. **C.** Immunoblot analysis of phospho-Chk1, phospho-H2AX, Sld5, and actin in cells overexpressing Sld5 or control constructs. UV-treated cells served as a positive control. Sld5 overexpression does not induce detectable DNA damage signaling. **D.** Immunoblot analysis of phospho-histone H3 in GL2 control and Sld5-depleted cells. Actin was used as a loading control. Sld5 depletion does not cause a marked increase in phospho-histone H3 under these conditions. **E.** Tumor proteomic analysis showing the association between GINS4/SLD5 and DYNC1H1 protein abundance across tumor samples. Samples are ordered by GINS4 expression and coloured by tumor group, supporting tumor-context-dependent co-variation between GINS4/SLD5 and DYNC1H1.

**Supplementary Fig. S4.**
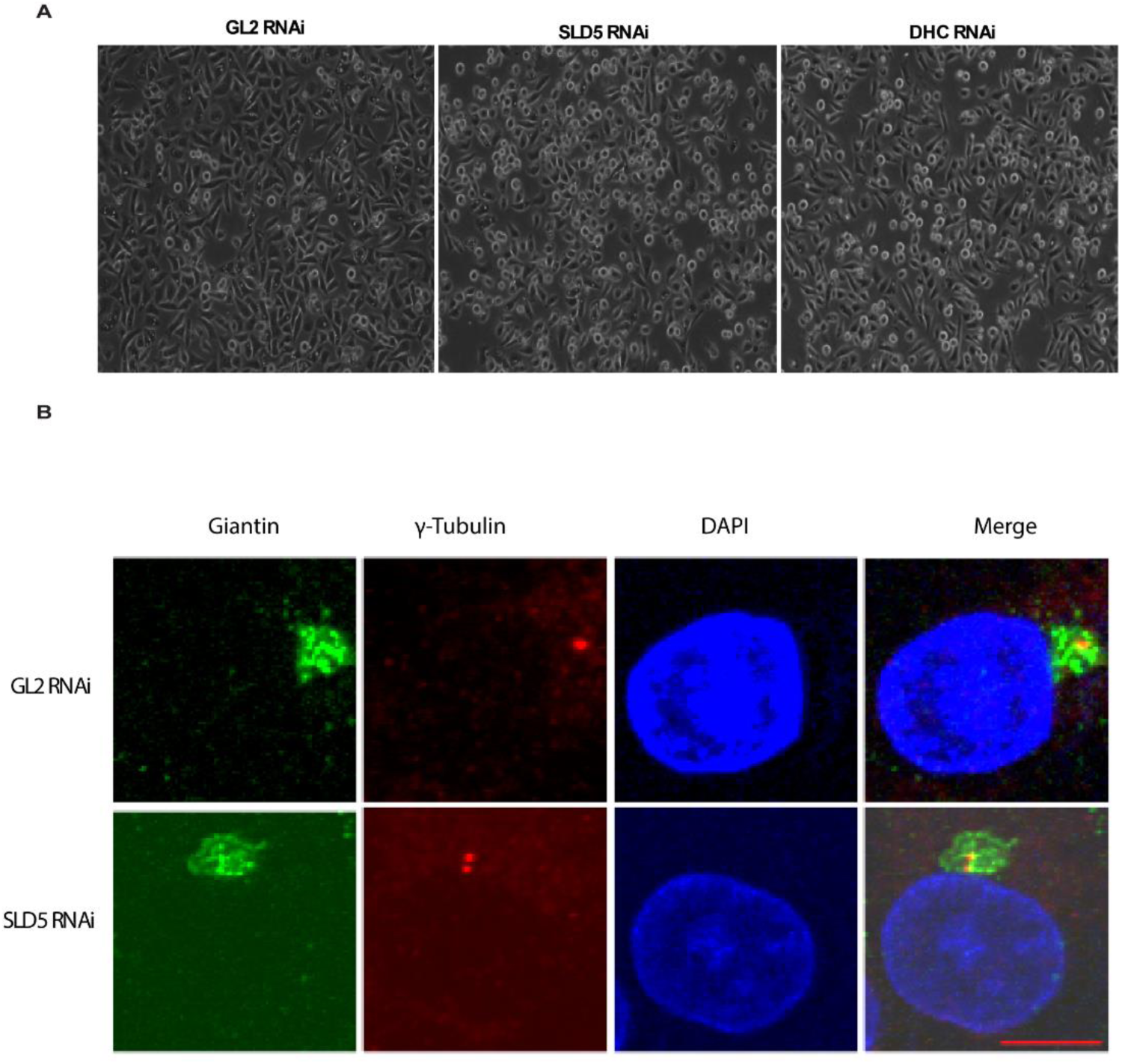
Sld5 depletion does not broadly disrupt Golgi organization. **A.** Representative phase-contrast images of GL2 control, Sld5-depleted, and DHC-depleted cells. Sld5 and DHC depletion produce comparable cellular perturbation, consistent with overlapping effects on mitotic and centrosome-associated processes. **B.** Representative immunofluorescence images showing localization of the Golgi marker giantin in GL2 control and Sld5-depleted cells. Cells were stained for giantin, γ-tubulin, and DAPI. Giantin retains a compact perinuclear Golgi-like distribution after Sld5 depletion, indicating that Sld5 loss does not cause a generalized disruption of organelle organization.

**Supplementary Fig. S5.**
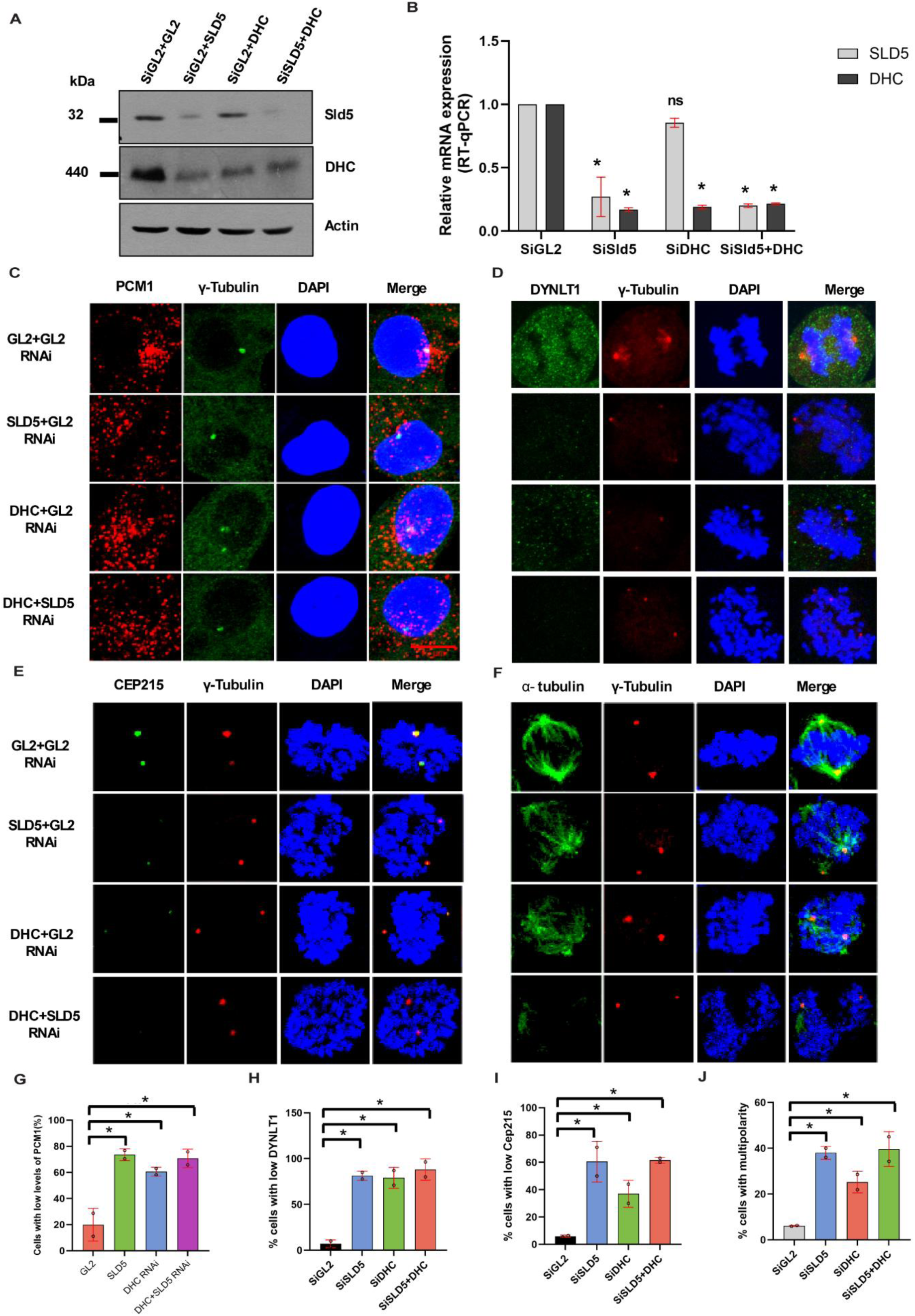
Co-depletion of Sld5 and DHC does not exacerbate centrosome–satellite defects, supporting a shared pathway. **A.** Immunoblot analysis of Sld5 and DHC protein levels after GL2 control RNAi, Sld5 RNAi, DHC RNAi, or combined Sld5 and DHC RNAi. Actin was used as a loading control. **B.** RT–qPCR analysis of SLD5 and DHC transcript levels after individual or combined depletion. Sld5 and DHC depletion is confirmed at the mRNA level. Bars show relative expression; error bars indicate variation among measurements. ns, not significant; *P < 0.05. **C.** Representative immunofluorescence images showing PCM1 localization after control, Sld5, DHC, or combined Sld5 and DHC RNAi. Cells were stained for PCM1, γ-tubulin, and DAPI. PCM1 is dispersed after Sld5 or DHC depletion, and combined depletion does not further increase the phenotype. **D.** Representative immunofluorescence images showing DYNLT1 localization after control, Sld5, DHC, or combined Sld5 and DHC RNAi. Cells were stained for DYNLT1, γ-tubulin, and DAPI. DYNLT1 signal is reduced after Sld5 or DHC depletion, with no clear additive effect after co-depletion. **E.** Representative immunofluorescence images showing CEP215 localization after control, Sld5, DHC, or combined Sld5 and DHC RNAi. Cells were stained for CEP215, γ-tubulin, and DAPI. CEP215 recruitment to centrosomes is reduced after Sld5 or DHC depletion. **F.** Representative immunofluorescence images showing spindle organization after control, Sld5, DHC, or combined Sld5 and DHC RNAi. Cells were stained for α-tubulin, γ-tubulin, and DAPI. Sld5 and DHC depletion produce spindle-pole defects, and combined depletion does not markedly exacerbate the phenotype. **G.** Quantification of cells with low PCM1 signal after individual or combined Sld5 and DHC depletion. **H.** Quantification of cells with low DYNLT1 signal after individual or combined Sld5 and DHC depletion. **I.** Quantification of cells with low CEP215 signal after individual or combined Sld5 and DHC depletion. **J.** Quantification of cells with multipolar spindles after individual or combined Sld5 and DHC depletion. Co-depletion does not yield an additive increase in multipolarity relative to the strongest single-depletion condition, supporting a shared Sld5–DHC pathway.

## Supplementary Table legends

**Supplementary Table S1. Antibodies list for immunofluorescence.**

**Supplementary Table S2. Antibodies list for immunoprecipitation.**

**Supplementary Table S3. Antibodies list for western blot.**

**Supplementary Table S4. Genes primer list for RT-qPCR.**

**Supplementary Table S5. Genome-wide Stouffer’s Z-score ranking from the integrated pan-cancer analysis.**

This table reports gene-level Stouffer’s Z-scores generated from the integrated pan-cancer analysis. Genes are ranked by their combined Z-score, with positive values indicating enrichment in the positive pan-cancer signal and negative values indicating enrichment in the opposite direction. GINS4 is listed within the positive tail of the distribution with a Stouffer’s Z-score of 9.1413. Columns include gene symbol and Stouffer’s Z-score.

**Supplementary Table S5A. H-score quantification of GINS4 immunohistochemical staining in representative normal and tumor tissues.** This table summarizes semi-quantitative H-score analysis of GINS4 immunohistochemistry in representative normal and tumor tissue sections. Normalized tumor staining was calculated relative to the indicated normal comparator. Liver cholangiocarcinoma showed a 2.67-fold increase in GINS4 staining compared with normal liver/hepatocytes, whereas prostate adenocarcinoma showed a 1.96-fold increase compared with the indicated normal comparator tissue. Columns include comparison, normal tissue, normal H-score, cancer tissue, cancer H-score, normalized normal value, normalized cancer value, and fold change.

**Supplementary Table S6. Cancer-lineage-specific association between SLD5 knockout effect and DYNC1H1 expression.** This table summarizes correlation analyses between SLD5 knockout-associated log fold-change and DYNC1H1 expression across cancer types. For each lineage, Pearson and Spearman correlations, coefficient of determination, regression slope, and regression intercept are reported. The analysis identifies tumor contexts with inverse or positive relationships between SLD5 loss and DYNC1H1 expression, supporting cancer-type-specific regulation of the SLD5–dynein heavy-chain axis.

**Supplementary Table S7. Cancer-lineage-specific association between SLD5 overexpression and DYNC1H1 expression.** This table summarizes correlation analyses between SLD5 expression in SLD5-overexpressing tumor contexts and DYNC1H1 expression across cancer types. For each lineage, Pearson and Spearman correlations, coefficient of determination, regression slope, and regression intercept are reported. Positive associations in several carcinoma lineages support coordinated regulation of GINS4/SLD5 and DYNC1H1 in selected tumor contexts, whereas weaker or inverse correlations indicate lineage-specific differences in this relationship.

**Supplementary Table S8. Pan-cancer correlation analysis of GINS4 and POLR2A expression.** This table summarizes cancer-lineage-specific correlations between GINS4/SLD5 and POLR2A expression. For each cancer type, Pearson and Spearman correlations, coefficient of determination, regression slope, and regression intercept are reported. Positive correlations across multiple tumor lineages support coordinated regulation of GINS4 and POLR2A, suggesting a potential connection between Sld5 and RNA polymerase II-associated transcriptional activity in cancer.

**Supplementary Table S9. Predicted kinase associations with GINS4/SLD5.** This table lists candidate kinases predicted to interact functionally with GINS4/SLD5 or to be associated with it. Genes are ranked by Z-score, with higher Z-scores indicating stronger predicted association. The top-ranked candidates include BUB1, WEE1, CDC7, EIF2AK1, VRK2, NME2, STK16, NEK2, TAF1, PLK4, TTK, and PLK1, highlighting mitotic checkpoint, centrosome maturation, and replication-associated kinase pathways as potential therapeutic nodes in SLD5-dependent tumors.

**Supplementary Table S10. Co-occurrence analysis of GINS4/SLD5-associated kinase candidates.** This table summarizes co-occurrence relationships among GINS4/SLD5 and candidate kinase genes. Columns report the number of cases with neither alteration, alteration of A only, alteration of B only, alteration of both genes, log₂ odds ratio, P value, q value, and co-occurrence tendency. Significant co-occurrence was observed among several kinase pairs, including GINS4–EIF2AK1, GINS4–WEE1, and GINS4–PLK1, supporting a coordinated kinase network linked to GINS4/SLD5-associated cancer dependency.

## Notes

### Competing Interest Statement

The authors have declared no competing interest.

